# Mobile genetic elements exhibit associated patterns of host range variation and sequence diversity within the gut microbiome of the European Honey bee

**DOI:** 10.1101/2025.01.31.635958

**Authors:** Chris R. P. Robinson, Adam G. Dolezal, Ivan Liachko, Irene L. G. Newton

## Abstract

Mobile genetic elements (MGEs), such as plasmids and bacteriophages, are major contributors to the ecology and evolution of host-associated microbes due to a continuum of symbiotic interactions and by mediating gene flow via horizontal gene transmission. However, while myriad studies have investigated relationships between MGEs and variation in fitness among microbial and eukaryotic hosts, few studies have incorporated this variation into the context of MGE evolution and ecology. Combining HiC-resolved metagenomics with the model honey bee worker gut microbiome, we show that the worker gut contains a dense, nested MGE community that exhibits a wide degree of host range variation among microbial hosts. Using measures of gene similarity and syntenty, we show that plasmids likely mediate gene flow between individual honey bee colonies, though these plasmids exhibit broad host range variation within their individual microbiomes. We further show that phage-microbe networks exhibit high variation among individual metagenomes, and that phages show broad host range with respect to both the number and phylogenetic distance of their hosts. Finally, we provide evidence that measures of nucleotide variation positively correlate with host range in bee-associated phages, and that functional targets of diversifying selection are partitioning differently between broad or narrow host range phages. Our work underscores the variability of MGE x microbial interactions within host-associated microbial communities and highlights the genomic variation associated with MGE host range diversity.

## 1 Introduction

Mobile genetic elements, or MGEs, are at the center of ongoing environmental and health-related crises due to their ability to transfer genes – include those conferring antibiotic resistance – via mechanisms of horizontal gene transmission (HGT). Plasmids and bacteriophages are among the most important important classes of MGEs due to their ability to mediate conspecific and interspecific HGT. These two classes are often embedded within dense microbial communities and exhibit high variation in transmission rate [1] and gene content [2], yet most microbial x MGE interactions are studied in homogenous, liquid cultures [3], which limits our understanding of the factors influencing MGE-encoded trait variability. The ecological and evolutionary impact of MGEs within natural microbial communities is far less understood. For example, phages can drive changes in microbial community composition [4, 5, 6, 7, 8], but evidence for this is inconsistent [9, 10, 11, 12]. Alternatively, the HGT-mediated transmission of genes often results in low genetic linkage among MGE-encoded genes. Variation in HGT rate among MGEs – and the genes they encode – likely plays a critical role in MGE evolution.

Recent work in several natural systems [13, 14, 15, 16, 5] have shown that viruses likely associate with several phylogenetically distant microbial hosts. Similarly, groups of highly similar plasmids have been shown to exhibit tremendous variation in host range [17]. The stability and frequency of these interactions are thought to be mediated by host population diversity, community diversity, and transmission rate [18, 19, 20, 21, 22]. In turn, variation at the level of MGE transmission rate has been shown to affect the evolution of phage virulence [23] and impact the gene content carried by plasmid replicons [24, 18]. However, MGE-associated host range variation has only been examined in a handful of systems, and our understanding of whether and how this variation might impact measures of genic diversity in MGEs remains an open question. A powerful model system with which to explore MGE dynamics within host-associated microbial communities is the Western Honey Bee (*Apis mellifera*) due to their long co-evolutionary history with their gut microbiome [25], which has resulted in a stable, relatively low complexity microbial community. A handful of studies have explored bee-associated MGE communities [26, 27, 28, 29, 30, 31] and have strictly focused on phages. This empirical work suggests that bees are associated with a highly diverse phage community that exhibit high variability in phage population sizes and rates of co-evolution with microbial hosts. CRISPR-based methods suggest that *Gilliamella, Lactobacillus*, and *Bombilactobacillus* are among the most frequently predicted hosts for bee-associated phages [26, 27, 28].

Here, we sought to further understand viral and plasmid dynamics within the honey bee worker gut microbiome. We hypothesized that both viruses and plasmids with variable host ranges will be prevalent within honey bee worker microbiomes. We further hypothesize that increasing phage host range will be associated with an increase in observed intrapopulation genomic diversity as a consequence of either co-evolutionary (such as diversifying selection on genes associated with microbial-phage interactions) or demographic processes (such as recent population expansion as phages encounter new hosts). To test these hypotheses, we conducted deep metagenomic and HiC-proximity ligation sequencing of three age-matched honey bee worker pools, each originating from one of three colonies governed by a single drone-inseminated queen and held in the same apiary. We show that colonies harbor similar microbial communities and plasmids (via relaxase and replicon typing) while phage communities are highly heterogenous and specific to each colony. We identify near-identical and syntenic plasmid-associated gene modules across all three honey bee colonies, suggesting either recent plasmid transmission or strong purifying selection on a subset of plasmid-associated genes. By examining HiC networks, we show that phage host range is dynamic within honey bee worker microbiomes, and that phage conspecifics exhibit wide variation in the number and phylogenetic divrsity of their microbial hosts. Finally, we show that increasing viral host range positively correlates with measures of genic variation. Using the honey bee model to study natural microbial-MGE dynamics *in vivo*, our results show that the gut microbiome contains a highly diverse MGE community that exhibits a wide range of host range variation among microbial hosts and highlight how this variation may affect both microbial and MGE evolution in a natural animal-associated microbial community.

## Results

### Recovery of viral, plasmid, and core bacterial phylotypes from the honey bee gut

We performed metagenomic and Hi-C sequencing of three pools of honey bee workers. Each pool consisted of dissected guts from 15 age-matched workers that were collected from one of three single-drone inseminated colonies at the Bee Research Facility at University of Illinois Champaign-Urbana. Microbes were enriched through differential centrifugation from these pools and used from gDNA extraction (see Methods). gDNA was used for metagenomic sequencing, yielding a total of 904,471,836 150-bp read pairs. The honey bee worker gut microbiome is dominated by 5 core microbial phylotypes [32], and here we recovered 12 high-quality (≥90% completeness and ≤10% contamination) and 1 mid-quality (≥50% completeness and ≤10% contamination) microbial genome by combining data from multiple binning strategies (see Methods). Microbial genomes were dereplicated at 97% ANI and used to recruit reads from all 3 metagenomes. 9 microbial metagenomically-assembled genomes (mMAGs) were retained following dereplication and are representative of 4 core phylotypes (*Giliamella apicola*, *Snodgrassella alvi*, *Bombilactobacillus mellifer*, and *Bombilactobacillus mellis*) and 5 non-core phylotypes (*Commensalibacter*, *Frischella*, *Enterobacter*, *Apilactobacillus*, and *Bartonella*) of the honey bee worker microbiome. Recovered mMAGs represent the majority of observed bacterial diversity in honey bee worker gut microbiome [33]. Across all samples, *Snodgrasella*, *Giliamella*, both *Bombilactobacillus* species, and *Frischella* phylotypes were dominant in terms of absolute genome-wide average read coverage (Figure 1A). Next, we characterized the presence and distribution of plasmids in the honey bee worker gut microbiome. Plasmids in honey bees are likely important mediators of functional traits through their associations with cargo genes (notably antimicrobial resistance cassettes) and as facilitators of horizontal gene transfer (HGT). Our investigation of the honey bee worker gut plasmidome led to the recovery of a total 41 plasmids from all metagenomes. Due to challenges inherent to plasmid assembly from short metagenomic reads and the high similarity of bacterial and plasmid-associated genes due to HGT, we used a previously benchmarked plasmid discovery pipeline [34]. First, plasmid sequences were called and clustered via MOB-suite [34], which uses a reference database containing sequences of known plasmid origin. We restricted our analysis to the recovery of these highly complete, high-confidence plasmids (pMAGs). Plasmid sequences within each metagenome were then dereplicated at 95% ANI across 85% of the plasmid length and against mMAG contigs. We note that while our approach is intended to reduce the instances of false positives in plasmid discovery, it likely results in a drastic under representation of the observed plasmid diversity in the honey bee worker gut metagenome. pMAGs varied greatly in size, exhibiting an average length of 9.7 kb (min. 2.2 kb to max. 55 kb). A minority of recovered pMAGs encoded genes for plasmid mobility, either through mobilization via another MGE (n = 11) or via conjugation (n = 1). pMAGs were classified via relaxase typing via MOB-suite, and several relaxase types were found in all three metagenomes (3 relaxase types) and in 2 of the 3 metagenomes (6 relaxase types). The remaining pMAG relaxase types were unique to each metagenome (Figure 1B).

**Figure 1:**
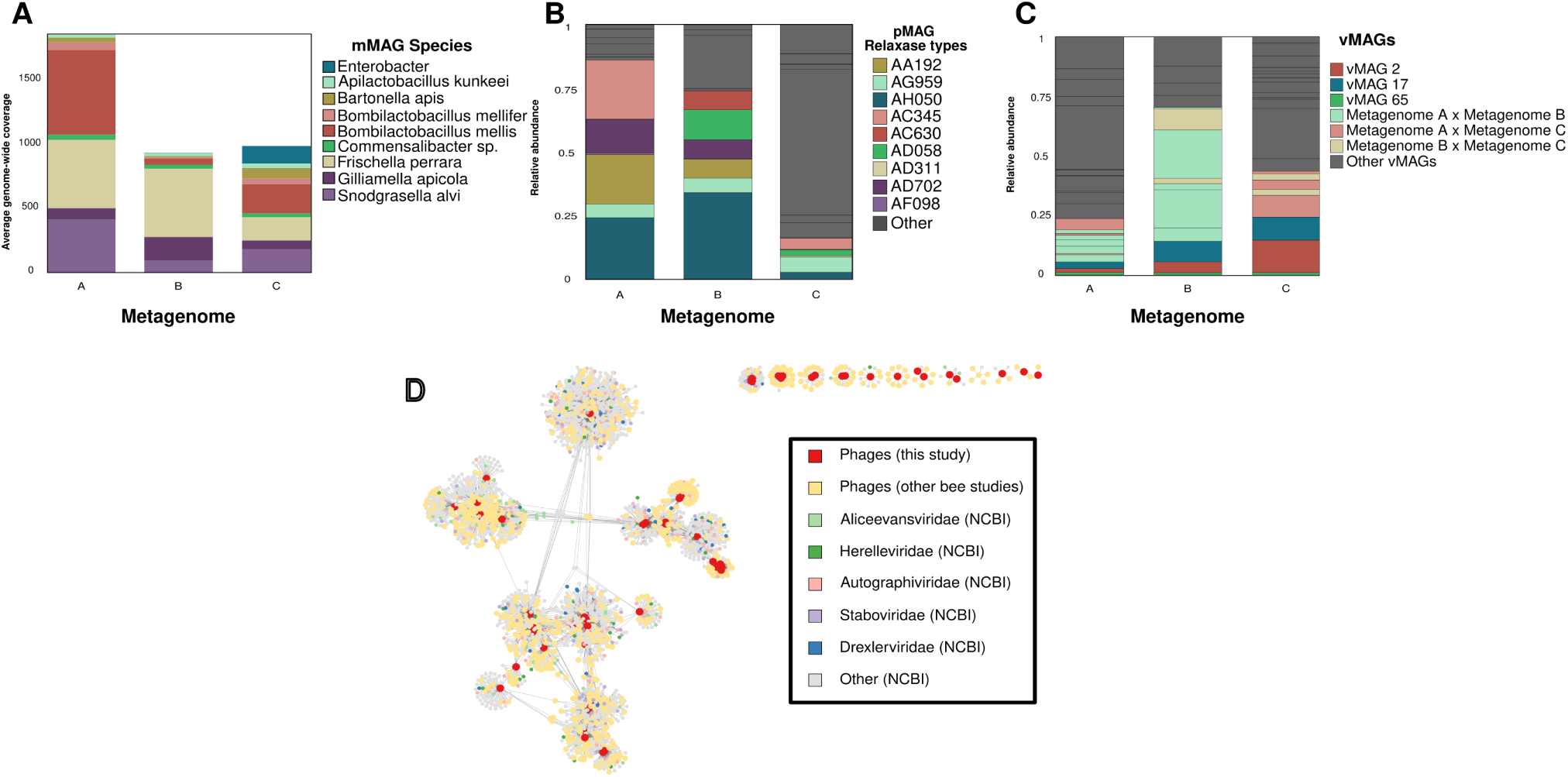
**(A)** Average genome-wide coverage for high-quality, dereplicated bacterial MAGs across all three metagenomes. **(B)** Relative abundance of plasmid bins associated with each metagenome. Colored portions represent the top 5 most abundant pMAGs across all samples. Shared colors are indicative of plasmids utilizing similar plasmid replicon types and are not representative of identical plasmids. **(C)** Relative abundance of viral MAGs across all samples. Colored portions represent abundance of vMAGs recovered from all three metagenomes. **(D)** vConTACT3 network generated from phage genomes recovered from this study (red), from all other bee phage studies where viral genomes were available (yellow) and from NCBI RefSeq (gray). 5 viral families exhibiting the most connectivity to bee phages are colored (Aliceevansviridae (light green), Herelleviridae (dark green), Autographiviridae (pink), Staboviridae (lavender), and Drexelviridae (blue).

In addition to pMAGs, we expanded our analysis to bacteriophages, leading to the recovery of 77 viral genomes that we could classify as complete (n = 8), high-quality (n = 27), or medium-quality (n = 42) via CheckV [35]. These genomes were dereplicated at 95% ANI and 85% breadth down to 70 genomes (vMAGs), which were used as the representative viral genomes for all downstream analyses. Our dereplication strategy reflects known species-equivalent OTU delineations in other bacteriophage communities [36, 37, 38, 39]. Dereplication resulted in a roughly 9% reduction in the number of viral genomes, suggesting a large degree of vMAG genomic variation across the three metagenomes. Similar to pMAG size, vMAG genome size was highly variable, averaging 28 kb (min 3 kb, max 143 kb) and encoding an average of 35 genes. A small subset (n = 5) of vMAGs were called as putatively proviral by CheckV. The relative abundance of vMAGs between metagenomes was highly heterogeneous (Figure 1C), exhibiting extensive variability relative to recovered pMAG relaxase types. To explore vMAG co-occurrence between colonies, we mapped reads from each metagenome to each vMAG and calculated the frequency of metagenomic variants associated with each vMAG by comparing the mean coverage of non-overlapping 100 bp windows to the mean intra-metagenomic vMAG genome coverage. A variant window was called if the mean coverage of the window was ≤ than 25% of the mean intra-genomic mean coverage for the associated vMAG. We used the frequency of these windows to determine the presence or absence of a vMAG within any metagenome other than the metagenome from which it was assembled (see Methods for full explanation). With this approach, we found that the majority of vMAGs were recovered from a single metagenome (n = 45). However, several vMAGs were identified as being shared across two (n = 22) or all 3 metagenomes (n = 3).

To further characterize the honey bee phage communities, we built gene-sharing networks to compare our vMAGs to previously described bee phage studies (Fig 1D, Supplemental Figure 1) [28, 29, 26, 27, 31], as well as reference phages taken from vConTACT3’s NCBI RefSeq database (v221). The majority of vMAGs from our analyses clustered with phage genomes described by other bee phage studies [31, 28, 29]. We assigned phage taxonomy to the family level in only 5.7% (4) of our 70 vMAGs. These vMAGs were identified as *Peduoviridae*. 30% of vMAGs (21) could not be identified at any taxonomic level while all other vMAGs were identified as *Caudoviricetes*.

### HiC-based linkage of mMAGs and MGEs

We next sought to understand how these MGEs are distributed among microbial hosts. Recovery of high-confidence linkages between hosts and MGEs, especially utilizing short-read metagenomic data, is difficult. Many approaches rely on matching CRISPR spacers, which are encoded in assembled microbial MAGs, to protospacers found in recovered MGEs. However, many bacteria do not use CRISPR systems [40], and the repetitive nature of CRISPR arrays make their assembly from short read metagenomic challenging. Further, naturally-recovered host-virus pairs have been increasingly shown to exhibit spatial and temporal variation [41, 42] in host range and fitness effects. This surprising heterogeneity has been observed between host bacterial populations [42], species [13], and even viral variants [41]. Within host-associated microbiomes, variation in viral host range is unlikely to be captured through the identification of historical interactions through spacer-to-protospacer matching via short-read metagenomic sequencing.

To address these challenges, we sequenced 3 Hi-C metagenomic libraries via the Phase Genomics ProxiMeta platform. After filtering, we recovered a total of 345,687,698 Hi-C paired-reads (104,694,834, 79,790,796, and 161,202,068 for metagenomes A, B, and C respectively). By mapping Hi-C reads to assembled mMAGs, pMAGs, and vMAGs, we can infer putative chromosomal-interactions between microbial genomes and intracellular MGEs. We estimated noise-to-signal ratios of raw Hi-C contacts using the ratio of intra-mMAG to inter-mMAG contacts (raw noise = 0.0343 ± 0.00417; noise-to-signal ratios were calculated as [13], see Methods for calculations of ratios.) Due to the possibility of horizontal gene transfer between MGEs (vMAG-to-vMAG, pMAG-to-pMAG, vMAG-to-pMAG), we restricted our analysis to include only contacts between mMAGs and MGEs (vMAGs or pMAGs). In the raw composite HiC network, we recovered 582,844 mMAG x MGE contacts while the number of mMAG x MGE contacts was reduced to 56,780 after normalization with HiCzin. In our composite network (Figure 2), we observed a high amount of variation in both the number of interacting contigs between any mMAG x MGE pair (visualized as thicker edges) and in the linkage strength (cumulative count of normalized contacts between mMAGs and MGEs; darker edges correspond to higher normalized linkage). Due to the wide variation in both the number of HiC connections connecting any mMAG x MGE pair and linkage strength, we calculated the average edge weight of HiC links between *Apilactobacillus kunkeii* and MGEs from each metagenome to use as the lower bound for which HiC links to include in our analysis. While *A. kunkeii* is widely distributed among honey bee larvae and the queen [30], its distribution and abundance in the worker midgut and hindgut is negligible, and we reasoned that many MGE x *A. kunkeii* contacts represent putatively spurious and weak interactions. Our results support this reasoning as *A. kunkeii* was the only mMAG that consistently aggregated both the weakest and fewest number of mMAG x MGE HiC links in both the composite and the individual networks (Supplemental Figures 2 - 5). After filtering, we saw a reduction of 7.85% in observed mMAG x MGE linkages for a total of 52,320 linkages in the composite network (Supplemental Figure 6). Because the HiC reads were mapped to a database built from all assembled high-quality mMAGs, pMAGs, and vMAGs, we performed a final filtering step to control for spurious contacts between highly similar genes shared by MGEs found in different metagenomes. Within each metagenome, we removed all mMAG x pMAG contacts from pMAGs that were not originally assembled from the same metagenome. Further, we used a stringent variant calling strategy to call the presence or absence of shared vMAGs across multiple metagenomes (see above section and Methods). mMAG x vMAG interactions from vMAGs called as absent within a metagenome were removed. The final composite network resulted in 9,289 high-confidence mMAG x MGE interactions.

**Figure 2:**
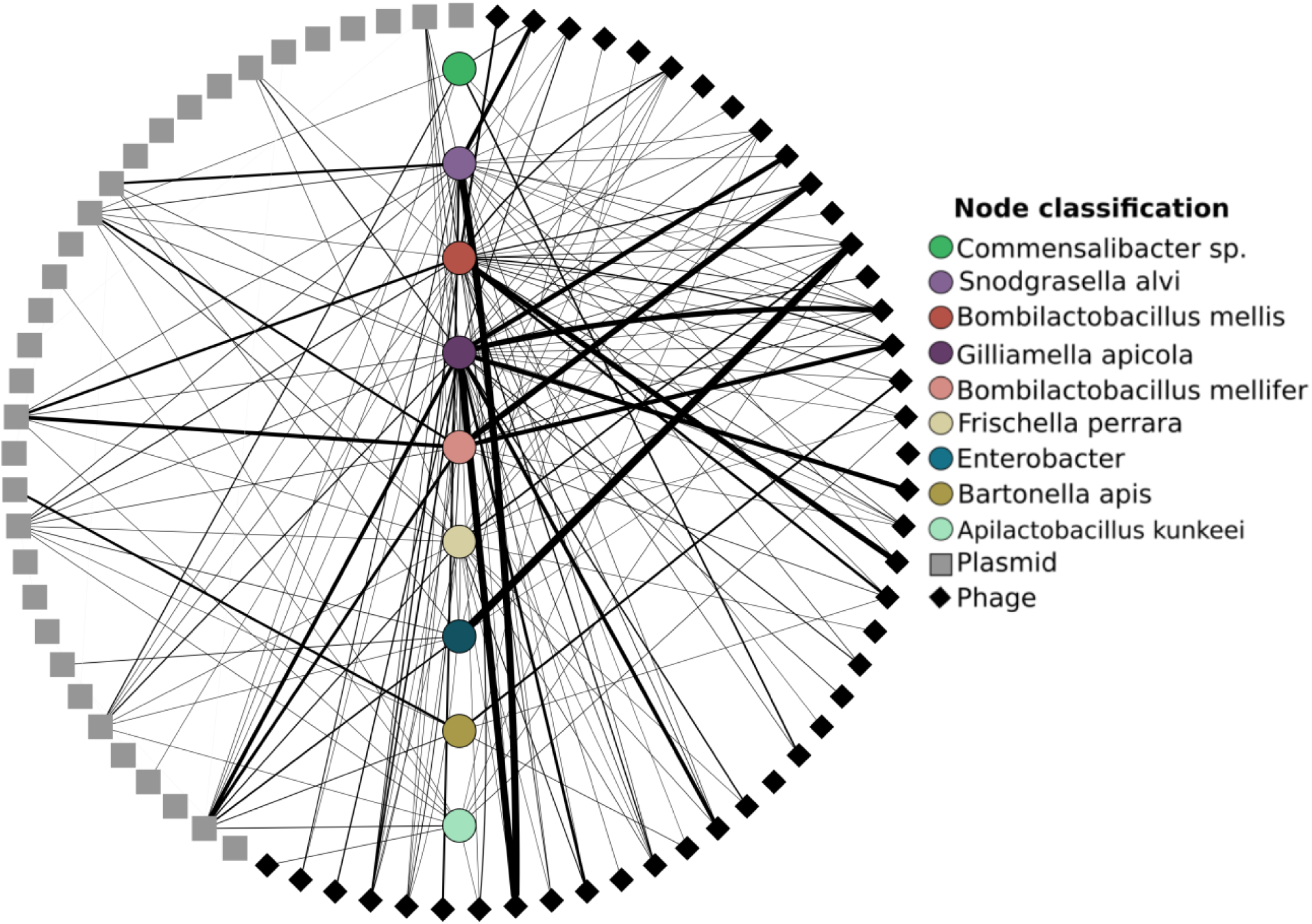
Cytoscape visualization of normalized HiC contacts between mMAGs, pMAGs, and vMAGs found in all metagenomes. mMAGs are centered in the circle and are colored by taxonomy. vMAGs are represented as black diamonds and pMAGs are represented as grey squares. Edge width corresponds to the number of contig-to-contig linkages between either vMAGs or pMAGs and mMAGs. The darkness of each edge corresponds to the cumulative strength of normalized HiC contacts between two node pairs.

We interpret darker, wider edges between mMAG x MGE pairs as indicative of the presence of intracellular associations between the pair, through we note that the presence of linkages between any mMAG x MGE pair is not proof of *in situ* association, nor is it proof of higher rates of infection within microbial populations. Rather, we use these signals to describe the distribution of MGE interactions across microbial hosts. Overall, *Snodgrasella*, *Giliamella*, and *Commensalibacter* aggregated the highest average number of mMAG x vMAG interactions (Supplemental Figure 4), while mMAG x pMAG interactions were driven mainly by *Commensalibacter*, *Snodgrasella*, and *B. mellis* (Supplemental Figure 2). 51.29% of mMAG x MGE linkages (4,764) were made up of mMAG x vMAG interactions while the remaining linkages (4,525) consisted of interactions between mMAGs and pMAGs. The viral linkages could be dereplicated down to 95 unique links shared between 9 mMAGs and 42 vMAGs while the pMAG x vMAG linkages could be dereplicated down to 53 unique links shared between 9 mMAGs and 12 pMAGs (Figure 2; individual metagenome networks are shown in Supplemental Figures 7 - 9). While the number of HiC connections between mMAGs and both classes of MGEs were relatively equal, we note biases intrinsic to HiC metagenome data. In particular, HiC signal is known to be biased towards higher coverage and longer contigs [43]. However, linear models showed no significant effect of read depth on the number of mMAG x vMAG HiC linkages (*p* = 0.8065, Supplemental Figure 10) or mMAG x pMAG HiC linkages (*p* = 0.8846m Supplemental Figure 11). However, larger MGEs (in terms of number of base pairs) were associated with differences in the number of HiC interactions. Larger pMAGs were associated with more HiC interactions (*p* = 0.0033, Supplemental Figure 12) while larger vMAGs were associated with fewer interactions (*p* = 1.79*e*^−12^, Supplemental Figure 13).

### Gene functions enriched in MGEs of honey bee workers

We identified a total of 2,228 and 655 genes on vMAGs and pMAGs respectively, which were then clustered into 18 protein clusters using KEGG BRITE KOs. These protein clusters can be broadly grouped into functions related to genetic information and processing, metabolism, and cellular signaling and overall represent a high degree of functional diversity. While both classes of MGEs were associated with a wide suite of systems relevant to their transmission and persistence (such as toxin-antitoxin systems, methyltransferases, and restriction-modification avoidance), we focused our analyses on MGE-encoded auxiliary metabolic genes (AMGs) (Supplemental Figure 14). Out of 591 recovered AMGs, 237 were associated with pMAGs, representing 36.2% of the total pMAG gene content. Antimicrobial resistance genes (AMR) were common and were recovered from 40.7% of all pMAGs in this study and from all individual metagenomes. These genes encoded functions related to resistance to tetracycline, chloramphenicol, and streptomycin. Genes encoding resistance to tetracycline were especially widespread on pMAGs, associating with several individual pMAGs in each of the individual metagenomes. These genes harbored similar organizational structure, encoding two *TetR* transcriptional regulators and a single MFS transporter (KF18476, K19047, and K08151). In contrast, vMAGs were associated a far fewer proportion of AMGs, encoding 111 AMGs which represented 4.98 % of the total genic diversity found on recovered phages. These viral AMGs were associated with a diverse set of functions, including iron efflux (K13283), fructoselyesine conversion (K19510), and domains associated with chlorhexidine efflux transporters (PF05232.16) which are part of a recently described set of antimicrobial efflux pumps known as PACE proteins [44]. Genes associated with lipopolysaccharide biosynthesis were found exclusively on vMAGs recovered in this study. vMAGs encoded 6 hypothetical chitinases (K03791), which were distributed among 5 unique vMAG genomes. A full list of MGE-associated annotations and their AMGs are provided as supplemental data (Supplemental Table 1).

### Near-identical gene modules found on plasmids in different honey bee colonies

While there was tremendous variation in HiC mMAG x MGE linkages within metagenomes, we reasoned that there may be additional gene-level variation at the level of MGE genomes as a consequence of MGE-mediated HGT. To identify genes associated with putative HGT events (i.e. mobile genes) from metagenomic short-reads, we used a previously benchmarked approach [45, 46] based on the identification of nearly identical genes present in phylogentically distant hosts. We used this method to identify 179 mobile genes from the 3 honey bee metagenomes. 63.5% of these mobile genes were associated with genes identified with pairs of pMAGs (Supplemental Table 2), while the remaining mobile genes were associated with genes identified on mMAG x mMAG pairs or vMAG x vMAG pairs. Plasmid-associated mobile genes were linked to plasmids recovered from all three metagenomes and represented 17.7% of all plasmid gene content, including 21 of the total 42 pMAG-associated AMR genes. In addition to AMR genes, we identified highly similar genes encoding functions such as transcription factors, transporters, chromosomal-associated proteins and prokaryotic defense systems between plasmid pairs. Many identified gene pairs were associated with plasmids with wide variation in host range as inferred by our HiC data. For example, pMAG AC 345 in Metagenome A is associated with no known detectable microbial hosts yet is associated with 3 putative HGT events to pMAGs recovered from the two other honey bee colonies (see Supplemental Figure 15 for Circos plot with identified pMAGs), which potentially expands the host range of these genes to 6 other bacterial hosts. Interestingly, plasmids encoding their own transmission machinery were no more likely to harbor mobile genes (Welch’s *t*-test, df=11.265, *p* = 0.09).

We next analyzed the synteny of nearly identical genes that were found on pMAG pairs by identifying pMAGs that contained ≥2 highly similar genes. Our goal was to understand the genomic context of how mobile genes were arranged on pMAGs as shared synteny would provide stronger evidence for gene mobilization. We identified 4 clusters of pMAGs containing syntenic genes exhibiting ≥ 99% similarity (Figure 3C). In cluster 1, comprising 46 mobile genes, we identified near-identical genes shared across multiple plasmid types and across multiple metagenomes. For example, plasmid AB595 from metagenome A, plasmid AB620 from metagenome B, and plasmid AD311 from metagenome C all contained a nearly identical (≥ 99%) gene pair (KEGG annotations: TetR/AcrR tetracycline repressor and MFS Transporter, tetracycline resistance protein). In this same cluster, plasmid AB595 from metagenome A is further connected to another pMAG in metagenome C (C plasmid AB190) via a set of paired genes encoding a transposon and phosphoryltransferase while C plasmid AB190 encodes a large antibiotic resistance cassette, encoding multiple genes for tetracycline resistance, transcription regulation, and DNA-binding. This entire antibiotic resistance cassette is nearly identical to cassettes recovered from pMAGs in the two other metagenomes (B plasmid AF098 and A plasmid AF098). Nearly identical cassettes are repeatedly observed on multiple pMAGs across all three metagenomes in all of the other clusters, including several genes encoding transmembrane secretion effectors [PF05977.17], ABC transporter domains [PF00005.31], and transcriptional regulations [PF21259.1] in cluster 3. In cluster 2, we observed several examples of AMR genes shared across multiple pMAGs in all three metagenomes. These AMR genes are flanked by several transposon genes, suggesting putative mechanisms of mobilization.

**Figure 3:**
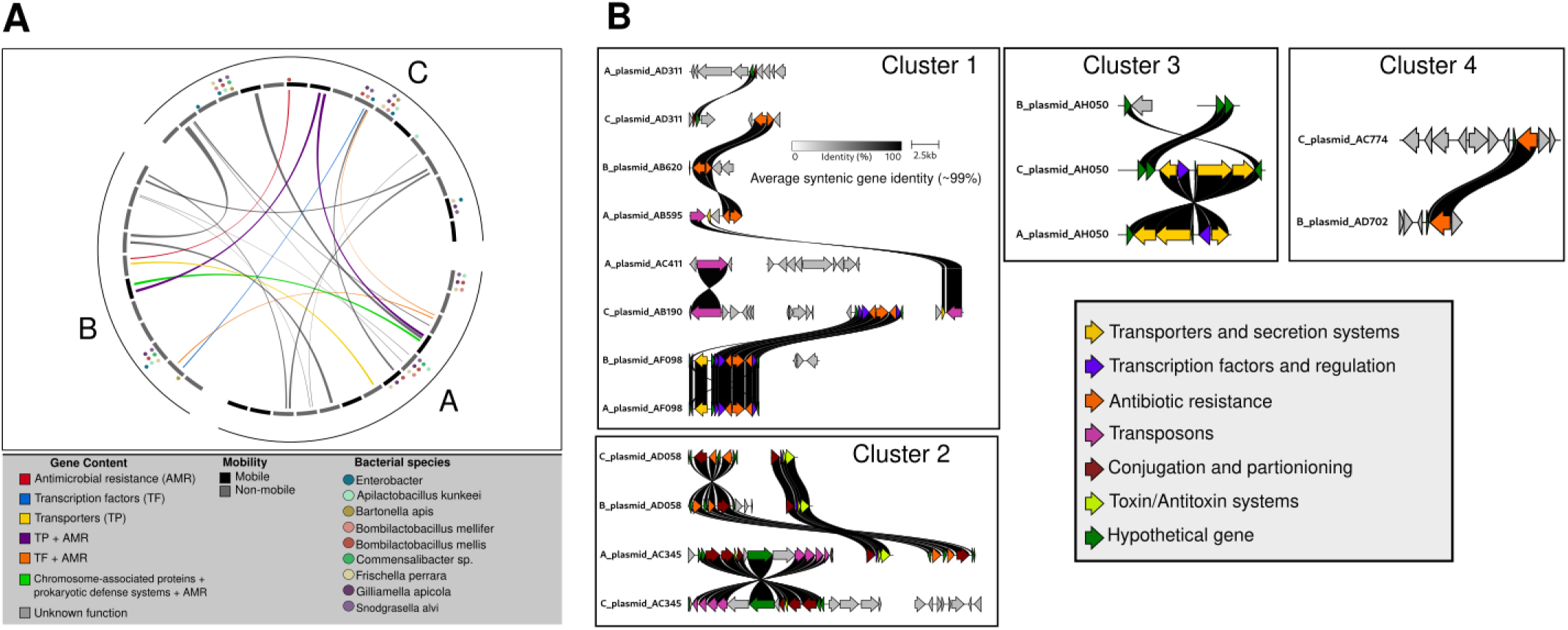
**(A)** Circos plot illustrating high similarity (≥97%) between genes found on all pMAGs recovered from each metagenome. pMAGs are demarcated by individual rectangles (inner arc) and are colored via the presence (black) or absence (grey) of genes encoding machinery for plasmid mobilization. Colored circles above each pMAG represent putative connections between pMAGs and mMAG hosts as inferred via HiC linkage data (Figure 2). All pMAGs found within a single metagenome are enclosed within a single arc (outer arc). Lines connecting pMAGs represent the presence of one or more genes with high similarity on both pMAGs. Lines are colored by the annotations associated with highly similar genes. **(B)** Gene arrow maps built from pMAGs sharing ≥2 genes of high identity. Arrows are colored by predicted function; Only ribbons corresponding to ≥99% identity shared between genes are shown. Metagenome identity for each pMAG is given as first character in pMAG ID (i.e, A plasmid AD311 was recovered from Metagenome A). Each cluster includes all pMAGs where at least one gene shared high similarity with another pMAG-associated gene.

### A subset of viral gene functions experience divergent patterns of natural selection

Due to the high observed heterogeneity within honey bee worker HiC networks, we next investigated whether differences in MGE host range exhibited any relationship with measures of genic diversity and adaptive evolution. We restricted our analysis to vMAGs for two reasons. 1) Our dataset includes a higher number of high-confidence viral genomes, and empirical data supports an ecologically and evolutionary-informed species-level delineation for metagenomically-assembled viral genomes [39]. 2) Phage genes are fairly well-studied and annotated relative to plasmid genes in our dataset which facilitates clearer interpretatability. To explore potential relationships between measures of genetic diversity and vMAG host range, we first calculated nucleotide divesity (*π*) for all viral genes in our dataset and compared the distributions of genic nucleotide diversity across four broad functional categories of viral genes (Figure 4A). Though we coarse-grained gene function by necessity, we found that genes encoding information processing functions harbored the least nucleotide diversity (mean 1.02*e*^−2^). Comparatively, phage structural genes exhibited large variation in nucleotide diversity (4.22*e*^−5^ to 9.96*e*^−2^, mean 1.39*e*^−2^) and included genes encoding functions related to phage capsid and tail assembly. Enzymatic and biosynthetic genes, as well as unannotated genes, were associated with similar measures of nucleotide diversity (mean of 1.749*e*^−2^ and 1.78*e*^−2^ respectively) and were significantly different from all other functional categories (Pairwise Wilcoxon Rank Sum Test, *p <* 0.05 (corrected for multiple comparisons). Bacteriophage holin genes, responsible for bacterial cell wall degradation, exhibited the highest levels of nucleotide diversity (mean of 7.85*e*^−2^) within genes related to biosynthetic and enzymatic function, and this may be a consequence co-evolutionary processes between phage and bacterial hosts [47, 48].

**Figure 4:**
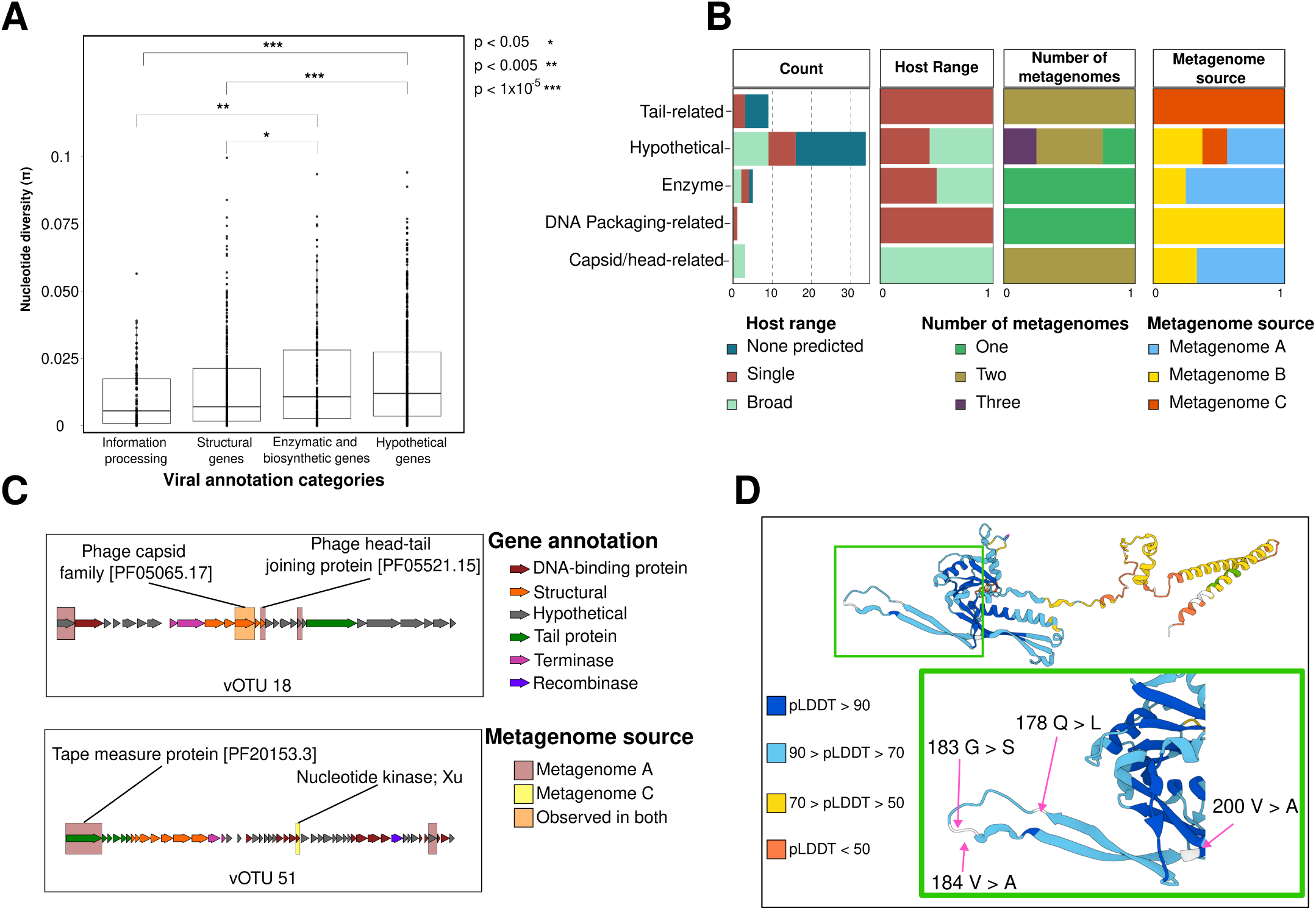
**(A)** Distributions of gene-wide nucleotide diversity *π* (y-axis) for viral annotation categories (x-axis). Significance differences between annotation categories were tested via Pairwise Wilcoxon Signed-Rank Tests. **(B)** Viral genes with pN/pS ratios ≥1 and their associated with viral metadata. Genes are grouped by gene function (y-axis) and their frequency within viral metadata (x-axis). Explored metadata includes the number of genes with pN/pS ratios ≥1, the number of unique HiC contacts associated with each vMAG; the number of metagenomes a given vMAG was recovered from; the metagenome that the vMAG was originally assembled from. **(C)** Gene arrow maps of select vMAGs. Arrows are colored by predicted function. Genes with pN/pS metagenomes 1 in metagenome A, metagenome B, or both metagenomes are higlighted in red, yellow, or orange respectively. **(D)** Alphafold protein prediction of Phage capsid family [PF05065.17] (see Fig 4C) that shows high similarity to the HK97 fold. Residues are colored by model confidence. Residues exhibiting high frequency (≥50%) intrapopulation polymorphism are highlighted in white. Amino acid variants are shown for specific residues on the protein’s E-loop (green box). Confidence values (pLDDT scores), taken from Alphafold, are provided.

We next sought to explore signatures of adaptive evolution in viral genes due to the differences in the distributions of *π* across categories of gene function. We calculated pN/pS ratios (the ratio of non-synonymous to synonymous polymorphisms) and identified 52 viral genes potentially undergoing diversifying or adaptive selection (pN/pS *>* 1); 23 of these genes were observed with pN/pS ratios *>* 2. 18 (34.5%) of these genes were associated with some annotated function, while the remaining 65.5% of genes were annotated as hypothetical. The largest class of annotated genes undergoing putatively diversifying selection were genes associated with viral tail proteins (n = 8), such as receptor binding tail proteins (vOTU 40) and tape measure proteins (vOTU 51). Genes encoding enzymatic proteins (n = 5) and capsid/head-related proteins (n = 4) made up the next two largest classes. Of the enzymatic genes with elevated pN/pS ratios, most predicted functions were related to nucleotide kinases, which may reflect selection for varying rates of DNA synthesis in replicating phage genomes [49] or in the regulation of phage gene networks [50]. Of the capsid/head-related genes associated with elevated pN/pS ratios, one gene had high similarity to the SPPI bacteriophage *gp16*, which as been shown to be involved in phage head-tail joining and facilitates the driving of phage DNA into host cells [51].

We further investigated the distribution of these genes based on vMAG metadata (Figure 4B). We found no clear association based on which metagenome the vMAG was recovered from nor the number of total metagenomes with which the vMAG was associated. Further, there was no evidence that genes with pN/pS ratios *>* 1 were enriched on phages associated with no mMAG hosts, a single host, or *>* 2 hosts (Pearson’s Chi-squared, *p* = 0.3). However, genes with pN/pS ratios *>* 1 and encoding tail-related proteins were strictly associated with vMAG genomes interacting with either zero or one mMAG species while all capsid/head-related genes that exhibited pN/pS ratios *>* 1 were found on vMAGs that interacted with ≥2 or more mMAG species.

Both tail-related and capsid/head-related genes with pN/pS ratios *>* 1 were recovered from vMAGs co-occurring in two metagenomes (Figure 4B). Since these metagenomic variants belong to the same viral species, we hypothesized that the same gene may be diversifying in both vMAG metagenomic variants as a consequence of similar selective backgrounds. Comparisons between metagenomic variants of the same vMAG was carried out by first identifying vMAGs with variants found in ≥2 metagenomes. These candidates were further screened by filtering out vMAGs that did not contain at least 1 gene with pNpS ratios *>* 1 in both variants. From our filtering strategy, we identified 2 vMAGs (vOTU 18 and vOTU 58) that were respectively associated with 4 and 2 genes that exhibited pN/PS ratios *>* 1 (Figure 4C). vOTU 18 co-occurred in metagenome A and metagenome C. In metagenome A, vOTU 18 was associated with 3 diversifying genes and exhibited 4 unique vMAG x mMAG interactions while in metagenome C, vOTU 18 associated with 1 diversifying gene and 2 unique vMAG x mMAG interactions. Likewise, vOTU 58 co-occurred in both metagenomes A and B and exhibited 1 unique vMAG x mMAG interaction in either metagenome, however all diversifying genes were found on the metagenome B-variant of vOTU 58. While most genes with elevated pNpS ratios were unique to metagenomic variants of both vMAGs, we identified a single gene (Phage capsid family [PF05065.17] that exhibited pNpS ratios *>*1 in both metagenomic variants of vOTU 18 (Figure 4C, highlighted in red). Using predictions from Alphafold [52], we found that the protein product of this gene was nearly identical to the well-characterized HK97 major capsid protein (or HK97 fold) (Figure 4D). After identifying the codon positions for each non-synonymous polymorphism in each metagenomic variant of vOTU 18, we mapped these positions onto the residues of our Alphafold model. We found several non-synonymous variants across the entire structure of the protein with variant frequencies exceeding 50% in both metagenomic variants of vOTU 18. However, we identified 5 non-synonymous variants in the metagenome C-variant of vOTU 18 that were located within the E-loop domain of the protein. At position 178, 62.1% of variants were associated with a glutamine residue while the remaining variants were associated with a leucine residue. The observed variation at this position results in changes in residue polarity, and likely affects the overall structure of the E-domain. HK97 major capsid proteins co-assemble with other capsid proteins to form a phage procapsid structure [53], and residue changes in this domain have been shown to affect the stability, size, and angle of the procapsid structure [53, 54].

### Increasing viral host range is positively correlated with measures of genomic variation

The distribution of genes with pN/pS ratios ≥ 1 with respect to variation in phage host range motivated us to more deeply explore relationships between host range and phage genic diversity. Here, we expand our definition of phage host range to include two alternative definitions of phage host range: intra-metagenomic host range and inter-metagenomic host range. We define intra-metagenomic host range as the number of unique vMAG x mMAG interactions within a metagenome while we define inter-metagenomic host range as the number of metagenomes that a single phage species is associated with. In both cases, we hypothesized that both intra-metagenomic and inter-metagenomic phage host range would positively correlate with higher values of allelic diversity (captured by *π*) and the number of SNVs (captured by *θ*w) due to a combination of changes in both selection regimes and/or demographic effects. Both *π* and *θ*w are estimators of a neutral mutation rate, *θ* = 2*Neµ* where *µ* is the population mutation rate and *Ne* is the effective population size. For each gene, we also calculated *Tajima’s D* (*D*), which can be understood as the difference between *π* and *θ*w.

Across all vMAGs, we recovered 1,375 genes with which we were able to calculate *π*, *θ*w, pN/pS ratios, and *D*. Mean genic values of *π* and *θ*w spanned as much as 3 orders of magnitude (Figure 5A) and are in line with previous estimates of these statistics using metagenomic data [55, 56]. Although mean read coverage values of *π*, and of *θ*w varied extensively between individual vMAGs and between vMAGs co-occurring across different metagenomes (Supplemental Figure 16), calculations of mean *π*, *θ*w, *D*, or pN/pS ratio values remained stable with respect to viral read coverage.

**Figure 5:**
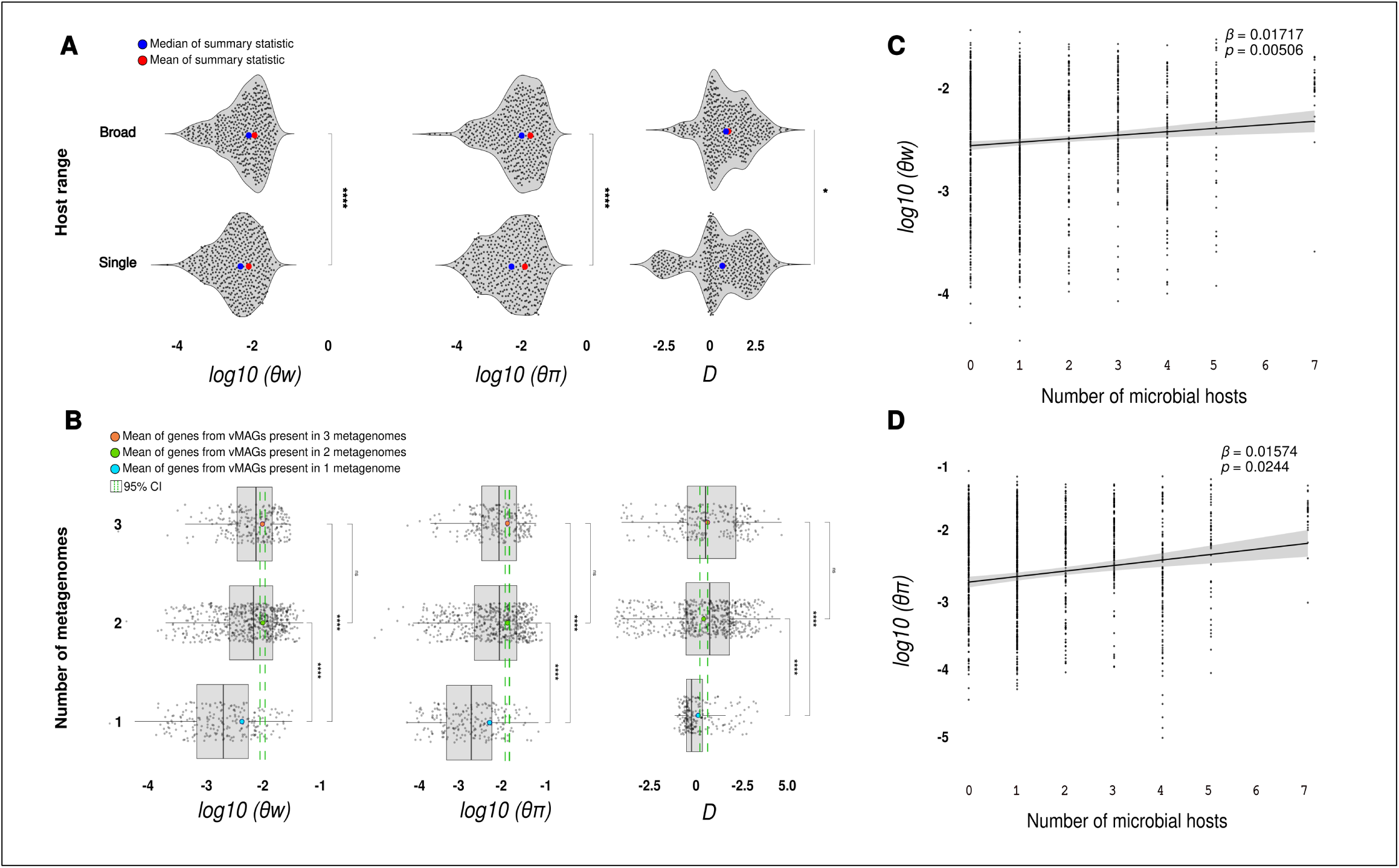
**(A)** Distributions of genic *θ*w, nucleotide diversity, and Tajima’s D (x-axis) from vMAGs associated with a single host (bottom distribution) or ≥2 hosts (top distribution). (**** = p *<* 10^−6^, * = p *<* 0.05). **(B)** Distributions of genic Tajima’s D, nucleotide diversity and *θ*w (x-axis) from vMAGs associated with 1, 2, or 3 metagenomes (y-axis). Gene-wide variation statistics are significantly higher in vMAGs associated with ≥2 metagenomes (*** = p *<* 10^−16^) **(C, D)** Correlations between the number of intrametagenomic mMAG x vMAG filtered HiC contacts (x-axis) and genic measures of *θ*w (C) and nucleotide diversity (*π*) (D). Regression line generated from geom smooth(method=lm) and shaded gray area represents 95% CI while p-values and beta coefficients were produced from linear mixed models.

Next, we compared differences in intra-metagenomic host range by comparing the distributions of genic *π*, *θ*w, and *D* values generated from narrow-host range vMAGS (single mMAG host) or broad range vMAGs (≥2 mMAG hosts). We found that the broad host range phages exhibited positive shifts in their distributions for all three summary statistics that were calculated (*π* (*p <* 1*e*^−9^); *θ*w (*p <* 1*e*^−7^); *D* (*p* = 0.04199) This trend continued when we compared differences in inter-metagenomic host range. We compared the distributions of genic *π*, *θ*w, and *D* values taken from vMAGs associated with 1, 2, or 3 metagenomes (Figure 5B, see Supplemental Figures 17 - 19 for metagenomic-specific viromes). All three distributions associated with broad inter-metagenomic host range vMAGs (present in ≥2 metagenomes) saw a significant positive shift when compared to vMAGs associated with only a single metagenome (Two-tailed Wilcoxon Rank Sum Test *p <* 1*e*^−16^). However, genes that were associated with vMAGs that occurred in 1, 2, or 3 metagenomes differened significantly in sample size. While the Wilcoxon Rank Sum test is fairly robust to unequal sample sizes, we addressed this inequality by comparing the mean of each summary statistic associated with vMAGs present in the smallest sample size (vMAGs present in 1 metagenome) to 95% confidence intervals produced from permutations of the largest distribution (vMAGs present in 2 metagenomes, see Methods). For each summary statistic, we found that the mean value of genes associated with vMAGs found in a single metagenome were completely excluded from the confidence intervals. The observed positive shift in estimators of *θ* for both intra- and inter-metagenomic host range suggests that viral host range expansion is associated with higher genic diversity. The positive shift of *D* in both cases is indicative of a higher number of intermediate-frequency polymorphisms, possibly as a consequence of balancing selection or recent population contraction.

Based on our analysis of viral functional genes, we reasoned that the positive shift in summary statistics could be caused by a number of confounding effects, such as annotation or the metagenomic origin of the vMAG. To investigate this further, we performed multiple regression analyses using linear models and linear mixed models to test for the correlation between intra-metagenomic host range and the summary statistics calculated above. In the linear mixed models, gene function, metagenomic source, and the number of metagneomes were treated as random effects. Using this regression approach, we found that intra-metagenomic host range exhibited a significant positive correlation with genic values of *π* (p *<* 6.4*e*^−6^, *R*^2^ = 0.03), *θ*w (p *<* 1.55*e*^−5^, *R*^2^ = 0.023), and *D* (*p* = 0.0042, *R*^2^ = 0.0794). Analyses using linear mixed models maintained these correlations for *θ*w (*p <*1*e*^−5^, Figure 5C) and *θ π* (*p* = 0.00208, Figure 5D). This positive correlation remained significant whether or not we included gene length and gene coverage in our models as a control for sequencing bias (*π* ~ intra-metagenomic host range (*p* = 0.0045); *θ*w ~ intra-metagenomic host range (*p <* 1*e*^−4^)). However, correlations between intra-metagenomic host range and measures of *D* were no longer significant (*p* = 0.142, see Supplemental Figure 20).

## Discussion

Mobile genetic elements drive a significant amount of adaptive evolution among microbial hosts within intra-host microbial communities, either through direct co-evolutionary interactions (anatagonistic co-evolution with phages) or through the acquisition of some ecologically important gene (such as plasmid-associated antibiotic resistance genes). However, interactions between MGEs and their microbial hosts are highly dynamic, and there is a severe paucity of information regarding microbe x MGE interactions *in situ.* Our study directly addresses this gap in knowledge by combining methods in metagenomics and proximity ligation (HiC) with the highly tractable honey bee worker as a model microbiome. Using conservative thresholds for MGE calling, we show that honey bee colonies co-housed in the same apiary exhibit highly variable phage and plasmid communities within age-matched worker guts and that these communities exhibit a large degree of genic and functional diversity. To our knowledge, our study is the first to formally investigate the plasmid component within honey bee worker metagenomic data. We used a rather conservative approach in order to minimize false positives arising from misassemblies and we look forward to further empirical work that explores plasmid ecology and evolution within the honey bee system. However, our observations of phage diversity recapitulate findings from other studies that investigate honey bee-associated phage communities [26, 27, 29, 31, 28]. We note that each of our metagenomic samples consists of 15 pooled workers, and our observations of MGE diversity may be driven by heterogeneity between individual worker microbiomes. The genetic background of workers has been shown to have an impact of strain-level variation within the worker microbiome [57], and we attempted to control for this through the use of single-drone inseminated queens, age-matched workers, and through sampling of hives co-housed in the same apiary, but future work is needed to characterize the community-level variation of MGEs between individual workers.

Through the use of HiC sequencing, we show that both viruses and plasmids likely interact with phylogenetically different microbial hosts in highly structured and extensively co-evolved communities, such as the *A. mellifera* worker microbiome. The observed host range variability of MGEs is likely an important factor in microbial evolution and ecology due to the high degree of MGE-encoded functional and genetic diversity. Indeed, while plasmids exhibited wide variation in host range, we found that over 17% of plasmid-encoded genes (and over 50% of ARGs) were nearly identical (≥97%) to a gene on another plasmid. We refrain from drawing strong conclusions regarding the presence of plasmid-mediated HGT in our study, as challenges in plasmid assembly from short-read data can contribute to misassembly and false positives. Alternatively, the high similarity of these genes across plasmids associated with different honey bee colonies may be due to the recent, independent introductions of these genes into honey bee colonies and/or strong-selective pressures within each individual colony. However, the conservation of gene synteny between plasmid pairs found in different colonies support recent plasmid dispersal and subsequent recombination as opposed to environmental selection on single genes. Understanding which mechanisms drive local adaptation within honey bee colonies – especially colonies proximal to one another – remains an open question [45]. As many of the observed plasmid gene-pairs may be transient, either due to neutral or deleterious fitness effects, future work should consider the impact of temporal and spatial variability on mobile gene dynamics, as patterns of mobile gene acquisition and stable co-occurrence may emerge from densely sampled colonies.

While broad host range plasmids were discovered nearly 50 years ago [58], the breadth and variation in observed phage host range here was surprising. Our observations are corroborated by several recent studies that use both HiC and CRISPR-based methods to resolve interactions between microbes and their bacteriophages [13, 14, 59]. Hwang et al [13] proposed several models to explain the observation of broad taxonomic host range in bacteriophages from a deep sea microbial mat, proposing scenarios such as high density of viral particles among a dense, highly diverse microbial community, the transfer of viral particles and/or genomes among syntrophic microbes, or bonafide host switching or host range expansion in viral populations. While our observations of genic variation in variants of the same viral species with respect to differences to viral host range might support the latter model of viral host switching or range expansion, we stress the need for further experimental work to validate the ability of virions to associate with phylogentically-distant hosts.

The observed patterns of molecular evolution based on population genetic summary statistics give further information about viral evolution with respect to host range variation. At least in the viral species that we analyzed, we found that phages that associate with a wider, more diverse set of hosts, are associated with a higher accumulation of genetic variation. The significant, positive shift in *Tajima’s D* associated with phages co-occurring in multiple metagenomes and in genes found on broad host range phages (Supplemental Figure 19) could be explained by the fixation of beneficial mutations on viral genes within some metagenomes but not others, balancing selection via interactions with multiple receptor proteins across bacterial species, recent population contraction in different populations of phage variants, or negative-frequency-dependent-selection as phage variants oscillate in frequency [60]. Regarding targets of selection, the divergent distribution of genes with high pN/pS ratios (*>*1) with respect to host range suggests a potential evolutionary consequence on phage gene evolution. However, we note that our calculations of *π* and *θ*w were made assuming an infinite sites mutational model, which assumes that mutations occur only once per site. Due to the high diversity that is observed in viral populations [39], our estimates of *π* and *θ*w are likely downwardly biased, though *π* is less affected than *θ*w [61]. Despite this bias, we interpret the positive correlations between intra-metagenomic host range and measures of nucleotide variation as a consequence of multiple non-exclusive scenarios. Though viral populations are likely very large, genetic drift is likely a major contributor in our system, as expansions in viral host range can drive differences in observed variation due to recent population expansion or bottlenecks. For example, the spread of some viral variant to a new microbial host (resulting in many low-frequency alleles) may co-occur with population contraction of another variant within the viral population. Alternatively, viral variants associating with a broader host range may be exposed to a diversity of selective regimes, and these regimes may not be homo-geneous across all variant subpopulations. While more work is needed to support these observations, they may provide clues with which to identify divergent genes in phages undergoing niche partitioning or understanding differences between specialist and generalist phages [62].

Interactions among mobile genetic elements and microbial hosts drive many fundamental evolutionary and ecological processes within host microbiomes. Developing a deeper understanding of these interactions is crucial if we are to generate testable hypotheses regarding the response of host-associated microbial communities to perturbation or disturbance, such as in the case of antibiotic usage. Our work highlights the variability of MGE x microbe interactions within host-associated microbial communities and suggests a deeper complexity in these systems than previously thought. Beyond the transfer of genes across wide phylogenetic distance and the driving of adaptive responses in microbial hosts, theory predicts that variation in HGT rates among plasmids affects plasmid gene gain and loss [63]. With respect to phages, many hypotheses that explore phage population dynamics, such as kill-the-winner or evolutionary arms races, often consider the co-evolution of a phage alongside a single bacterial host species. While we await further empirical validation of phage host range variation [64], the observations of this study (and others [13, 14, 58] suggest that bacteriophages may be engaging in antagonistic co-evolution among multiple bacterial species within a community. Future ecological and evolutionary work may benefit from incorporating these observations into predictive models.

## Methods

Detailed protocols are available in SI Appendix, Supplementary Methods

### Sampling, metagenomic assembly, and HiC reconstruction

Samples (N = 3) were collected from managed honey bee colonies at the University of Illinois Champaign-Urbana in 2022. Samples were composed of 15 pooled worker guts (in-hive bees) which were homogenized, flash frozen, and sent for gDNA extraction, short-read metagenomic sequencing, and HiC proximity ligation via Phase Genomics’ ProxiMeta service. All libraries were sequenced on a single lane of an Illumina Novaseq. Microbial, viral, and plasmid sequences were called from each metagenome, dereplicated, and annotated using a bespoke bioinformatic pipeline. Only high-confidence, dereplicated assemblies were retained. We performed a pairwise BLAST to identify nearly identical genes found on MGEs assembled from different individual metagenomes. To determine bacteriophage co-occurrence across metagenomes, we used a conservative read recruitment strategy to identify regions of low coverage. Phages exhibiting few regions of low coverage (*<*25% of genome length) were considered to be co-occurring in multiple colonies. We generated a Hi-C network from MAGs using the conctact matrices generated from the Hi-C proximity ligation sequencing data. These matrices were normalized using HiCZin [65]. Contacts between MGEs and *Apilactobacillus kunkeii* were used as a lower bound to remove spurious interactions.

### Calculation and inference of measures of genic variation

Metagenomic reads were mapped to vMAGs using bowtie2 (–sensitive-mode). Sorted BAM files and the vMAG contigs were used as input for Anvi’o [66] which was used to call single nucleotide variations across viral contigs. Microdiversity metrics (nucleotide diversity, Watterson’s theta, Tajima’s D, and pN/pS ratios) were calculated from SNVs using custom Python scripts. We tested for differences among these summary statistics across different groups of gene function using two-tailed Wilcoxon Rank Sum tests. Genic variation among viruses co-occurring among one, two, or three metagenomes was tested via two-tailed Wilcoxon Rank Sum tests and permutation testing to account for differences in sample size among viral groups. Relationships between summary statistics and variation in viral host range was examined via two-tailed Wilcoxon Rank Sum tests and linear-mixed models.

## Acknowledgments

We thank Audrey Parish for the sampling of honey bee colonies and for sending the pooled worker tissue to Phase Genomics. We thank Moutusee Islam for the preparation and sequencing of the metagenomic and HiC libraries and Tara Rickman for data interpretation and initial visualization of HiC data. We are grateful to Lílian Caesar and L. Felipe Benites for their help in data analysis and feedback on the writing of the manuscript. This work was financially supported by a Costco/Project Apis m. research grant to C.R.P.R., an NSF IOS Collaborative Research award (2005306) and by an NSF DBI Biology Integration Institutes award (2022049) to I.L.G.N. This work was partially supported through a grant from the Bill and Melinda Gates Foundation to Phase Genomics.

## Data availability

Raw metagenomic and HiC reads are available from NCBI under BioProject PRJNA1206451. Links to many of the Python workflows, as well as greater detail regarding how our analyses were done, can be found here: (https://github.com/en-nui/HoneyBeeHiC_public).

## Notes

### Competing Interest Statement

The authors have declared no competing interest.

