## Supplemental Information for "Mobile genetic elements exhibit associated patterns of host range variation and sequence diversity within the gut microbiome of the European Honey bee"

**This PDF file includes:**

Supplementary Methods

Supplemental Figures and legends S1 to S20

References for supplemental material

### Supplementary Methods

#### Honey bee management, metagenomic sequencing, and HiC sequencing

Honey bee samples were collected from managed single-drone inseminated (SDI) colonies at the Bee Research Facility at the University of Illinois Champaign-Urbana. SDIs were chosen to minimize intra-colony variation among workers [1]. Each colony was located within the same apiary in order to minimize variability associated due to changes in distance and acquired environmental resources. For each metagenomic sample, a frame was pulled from the colony and visually inspected for the queen. Workers were quickly brushed from the frame and moved to 4C until bees were immobilized. For each individual colony, 15 age-matched workers were removed and dissected, retaining the entire gut of the worker. These guts were pooled and transferred to a sterile dounce homogenizer. 3 ml of cold PBS was added to each homogenizer and the bee tissue was homogenized on ice. 3 ml of homogenate was then pipetted into 3 sterile 1.5 ml microcentrifuge tubes and spun down at 500g for 10 minutes to pellet host tissue. Supernatant from all 3 microcentrifuge tubes was transferred to a sterile Falcon tube. 1.5 ml of the pooled supernatant was then transferred to a new microcentrifuge tube and spun down at 5000g for 10 minutes. Following this last centrifuging step, the supernatant was removed and the bacterial pellet was flash frozen with liquid nitrogen before being stored at -80C. This process was repeated for all metagenomic samples. All samples were sent to Phase Genomics for DNA extraction and the generation of short read and Hi-C proximity ligation libraries via Phase Genomics' ProxiMeta service. Hi-C libraries were prepared using the restriction enzymes, Sau3AI and MluCI. Both Hi-C and 150 bp short read libraries were sequenced on a single lane of an Illumina NovaSeq.

#### mMAG assembly, binning, and annotation

Reads from the metagenomic library were filtered by quality score and trimmed using BBduk (BBMap - Bushnell B. - sourceforge.net/projects/bbmap/) before assembly with MEGAHIT [2] (–min-contig-len 1000 –k-min 21 –k-max 141 –k-step 12 –merge-level 20,0.95). Microbial metagenomically-assembled genomes (mMAGs) were binned by combining results from multiple binning software, including Maxbin2 [3], MetaBAT2 [4], and HiCBin [5]. These bins were used as input for DASTool [6]. mMAG quality was evaluated using checkM [7]. We retained only medium-to-high-quality mMAGs ($\geq$ 50% completeness and $\leq$10% contamination) from each metagenome. mMAGs from each metagenome were dereplicated at 97% average nucleotide identity (ANI) using dREP [8]. mMAGs were taxonomically assigned using GTDB-Tk [9] on KBase [10]. Dereplicated mMAGs that were representative of the known phylogenetic diversity within honey bee workers were used for all further analyses. mMAG genes were predicted using Prokka [11] and annotated using DRAM [12] and METABOLIC [13] using default databases (DRAM-setup.py prepare_databases –output_dir DRAM_data –skip_uniref).

#### vMAG binning and annotation

Contigs from each metagenome and $\geq$ 1000 bp in length were predicted as viral using VIBRANT v1.2.0 [14]. Contigs predicted as viral by VIBRANT were used as input for CheckV [15]. Viral contigs that were classified as medium to high-quality by CheckV were retained for all further analysis. Retained viral contigs were dereplicated with CD-HIT [16] at 95% ANI and 85% breadth (cd-hit-est -i $cat_file -o $cat_file_replicate -c 0.95 -n 10 -aS 0.85). Dereplicated viral contigs (vMAGs) were taxonomically identified using vConTACT3 (https://bitbucket.org/MAVERICLab/vcontact3/src/master/). vMAGs were annotated using Cenote-Taker2 [17] and DRAM-V and were screened for putative auxiliary metabolic genes (AMGs) using both VIBRANT and DRAM-V.

#### pMAG binning and annotation

Plasmid contigs from each metagenome were predicted using MOB-recon, a subprogram of MOB-suite [18]. Contigs $\geq$ 1000bp in length in MOB-recon. Predicted plasmids were then clustered within each metagenome at 95% ANI and 85% breadth using CD-HIT. Due to the difficulty in assembling plasmids from short-read metagenomic datasets, all plasmid bins were checked for contigs that had assembled into both mMAG and plasmid bins. These contigs were excluded from the plasmid bins. Finally, all plasmid bins from all metagenomes were dereplicated at 95% ANI and 85% breadth to allow for comparison between metagenomes. These dereplicated plasmid bins (pMAGs) were annotated using DRAM and METABOLIC. Because DRAM and DRAM-V expects either microbial or viral contigs, we could not utilize AMG scoring for pMAGs. Instead, pMAGs were manually inspected for the presence or absence of AMGs via annotations derived from KEGG [19] and PFAM [20].

#### HiC-based reconstruction of mMAG and MGE associations

Hi-C reconstruction of mMAGs and MGEs closely followed methods described by Hwang et al. [21]. Hi-C reads were quality filtered using BBduk and mapped using bwa mem (bwa mem -5SP) [22] against a combined database containing all mMAGs, pMAGs, and vMAGs. All contigs in this database were dereplicated a final time using CD-HIT at 95% ANI and 85% breadth to minimize Hi-C reads matching across highly similar contigs. Metagenomic short reads from each individual metagenome were also mapped against this database and read coverage was calculated using bbduk. Hi-C contact maps were normalized using the unlabeled version of HiCZin [5] (hiczin.py norm -e Sau3AI -e MluCI). Normalized contacts were further filtered by removing all contacts that exhibited less contact strength (e.g. HiC paired read coverage) than the average contact strength between *Apilactobacillus kunkeei* and *A. kunkeei*-associated MGEs. Hi-C based contacts between mMAGs and MGEs were visualized using Cytoscape 3.10.2 [23].

#### Inference of horizontal gene transfer between mMAGs and MGEs

To identify putative horizontal gene transfer events, we used a method benchmarked by Smilie et al. [24] and Brito et al. [25]. All contigs in the deduplicated database described in the previous section (HiC-based reconstruction of mMAG and MGE association) were screened in a pairwise manner with BLAST (blastn -query $f -db $db -evalue 1e-6 -perc_identity 97 -outfmt '10'). Previous approaches used an identity threshold of ≥99% sequence identity with sequences ≥500 bp in length between any two distantly related genomes [25]. We relaxed this threshold to ≥97% as previous work [26] found that 16S rRNA gene sequences from distinct core phylotypes in the honey bee microbiome cluster at ≥97% sequence identity. We note that identification of highly identical sequences using this method does not imply direct transfer between any plasmids.

#### vMAG population genomics and analyses

Filtered reads from each metagenome were mapped to all conserved single-contig vMAGs using bowtie2 [27] in sensitive mode. We calculated the average genome-wide coverage for each vMAG using bedtools [28]. vMAG contigs, read coverage data, and BAM files were given as input for Anvi'o [29]. Anvi'o was used to call single nucleotide variants (SNVs) across all viral contigs. SNVs were used to calculate microdiversity metrics using custom Python scripts. These scripts are available here: (https://github.com/en-nui/HoneyBeeHiC/blob/main/python_home/popgen_annotations_summary_statistics_program.py

#### Generation of pN/pS ratios

The pN/pS ratio is the ratio of intra-population non-synonymous (pN) and synonymous (pS) rates. It is similar to estimations of the dN/dS ratio which can be compared across different bacterial strains or species. Analogous to interpretations of dN/dS ratios in protein-coding genes, we can detect putative instances of purifying selection (pN/pS $< 1$), neutral evolution (pN/pS $\approx 1$), and positive selection (pN/pS $> 1$). pN/pS ratios for each viral protein-coding gene were estimated via Anvi'o (*anvi-get-pn-ps-ratio*) [29, 30].

#### Calculation of vMAG nucleotide diversity

Microdiversity parameters for viral genes were calculated at the gene and genome-wide level. We calculated nucleotide diversity ($\pi$), Watterson's estimator ($\theta_w$), and *Tajima's D* [31]. All statistics were calculated using assuming an infinite sites mutational model. Nucleotide diversity ($\pi$), which measures the average number of pairwise nucleotide differences between any two sequences [32] was calculated from SNVs called by Anvi'o via metagenomic reads mapped to viral contigs as follows:

$$\pi = \frac{\sum_{i<j} k_{ij}}{n(n-1)/2}$$

where $k_{ij}$ equals the number of nucleotide differences between the $i$th and $j$th sequences in the sample and the denominator represents the number of unique comparisons made between $n$ sequences [32]. $\theta_w$, which is an alternative estimator of $\theta$, was calculated as follows where $S$ is equal to the total number of segregating sites (or SNVs):

$$\theta_w = \frac{S}{a}$$

where $a$ (a normalizing factor representing the sample size (n) is calculated from:

$$a = \sum_{i=1}^{n-1} \frac{1}{i}$$

Because both $\pi$ and $\theta_w$ are both estimators of the same parameter $\theta$, the expected difference between them should be 0 under the standard neutral model. We estimated the differences between $\pi$ and $\theta_w$ via *Tajima's D* [31] as follows:

$$D = \frac{\pi - \theta_w}{\sqrt{Var(\pi - \theta_w)}}$$

#### Statistical analyses

All statistical analyses were performed using R v4.4.1. Prior to building linear regressions and linear mixed models, all data was standardized using the Box-Cox transformation in order to ensure the normality of residuals. All linear regression analyses and *t*-tests were built using the *R* function, "summary.lm()" [33].

#### Noise-to-signal calculations

Raw noise-to-signal ratios were calculated as Hwang et al. [21]

$$\frac{Number\ of\ inter\text{-}mMAG\ HiC\ contacts}{Number\ of\ intra\text{-}mMAG\ HiC\ contacts}$$

where *inter-mMAG contacts* is equal to the number of HiC read pairs mapping to different mMAGs and the number of *intra-mMAG contacts* is equal to the number of HiC read pairs mapping to the same mMAG.

#### Identifying vMAG species across metagenomes

To identify if a vMAG species was associated with 2 or more metagenomes, we mapped reads from each metagenome to a database of all recovered, dereplicated vMAGs. Read depth files were generated for each vMAG-metagenome pair. To account for viral variation due to recombination and HGT, we utilized the frequency of viral metagenomic islands (MGI) within each metagenome to determine the presence or absence of a vMAG variant. First, mean genomic coverage was generated for each vMAG-metagenome pair. Pairs with mean coverages $\leq$ 5x were removed. Next, each pair was split into non-overlapping 100-bp windows. The mean coverage was calculated for each window, and the window was flagged as an MGI if the mean coverage of the window was less than 25% of the mean coverage for the vMAG genome. A vMAG was considered to be in multiple metagenomes if the sum length of the metagenomic islands was greater than or equal to the 25% of the total vMAG genome length.

#### Effect of read depth, contig length, and MGE size on HiC networks

The effect of read depth and MGE size on the number of mMAG x MGEs HiC linkages was tested via multiple linear regressions:

$$Y \sim \mathit{Read\ depth}$$

$$Y \sim \mathit{MGE\ size}$$

$$Y \sim \mathit{Contig\ length}$$

where Y represents either the number of HiC contacts or the unique number of microbial hosts for either vMAGs or pMAGs. The significance of these multiple linear regressions were evalulated with the $t$-test of the R function *summary.lm()*. The *p.adjust()* function of the R package "stats" was used to adjust the linear regression $P$-values using FDR correction.

#### 0.1 Variation across annotation categories and presence in multiple metagenomes

Measures of genomic variation ($\pi$, $\theta$w, and Tajima's $D$) were calculated separately for each vMAG-associated gene and binned into one of 4 major annotation categories (information processing, structural genes, enzymatic and biosynthetic genes, and hypothetical genes). These categories were built from KEGG BRITE Pathway categories. Differences in genomic variation among the 4 functional groups were tested via Pairwise Wilcoxon Rank Sum tests. $P$-values were corrected for multiple tests as above. This procedure was applied to test for differences in genic variation among vMAGs associated with 1, 2, or 3 metagenomes. Plots were generated with ggplot2 [34].

#### Effect of host range on viral gene sequence variation

To investigate the relationship between viral host range and measures of genic variation, we used linear mixed models through the R package, lme4 [35]. We opted for linear mixed models to better incorporate random effects. Random effects are grouping factors that explain random variance of the relationship between the response variable and fixed effects across a number of different groups [36]. Further, our data violates assumption of independence due to the presence of vMAGs in 2 or more metagenomes as well as the possibility of HGT-mediated acquisition of near-identical genes across vMAGs. Linear mixed models are robust to these violations. Before regression modeling, we removed all vMAGs with 0 vMAG x mMAG interactions across all metagenomes. Next, data was normalized using the Box-Cox transformation to ensure the residual normality. The regression models are presented as below:

$$\theta\pi^{*} \sim \textit{Viral host range} + \textit{Annotation} + \textit{Metagenome source} + \textit{Number of metagenomes}$$

$$\theta w^{*} \sim \textit{Viral host range} + \textit{Annotation} + \textit{Metagenome source} + \textit{Number of metagenomes}$$

$$\textit{Tajima's } D^{*} \sim \textit{Viral host range} + \textit{Annotation} + \textit{Metagenome source} + \textit{Number of metagenomes}$$

where $\theta\pi$, $\theta$w, and *Tajima's D* are values calculated for each vMAG-associated gene (n = 1375). Intra-metagenomic host range was defined as the number of unique intra-metagenomic mMAG hosts. Annotation, metagenome source, and number of metagenomes were included as random effects to account for sequence variation between these groups. Asterisked values incorporate gene coverage and length into the model as random effects. *P*-values and effect coefficients for each regression model was calculated via the R package "lmerTest". Tests of differences in the measures of genic variation associated with differences inter-metagenomic host range (defined as the number of metagenomes with the same co-occurring viral species) was done via two-tailed Wilcoxon rank sum tests on untransformed data. To account for differences in sample size associated with each group, *N* number of genes were sampled from the larger distributions, where *N* is equal to the number of genes in the smallest distribution (vMAG genes present in a single metagenome). This process was repeated 1000 times to produce a 95% confidence interval around the mean of the larger distributions. *P-values* were calculated for both the Wilcoxon Rank Sum Test and from the frequency with which the mean of the single-metagenome vMAG genes was found within this 95% confidence interval. All plots were generated with ggplot2.

### Supplemental Figures S1 to S20

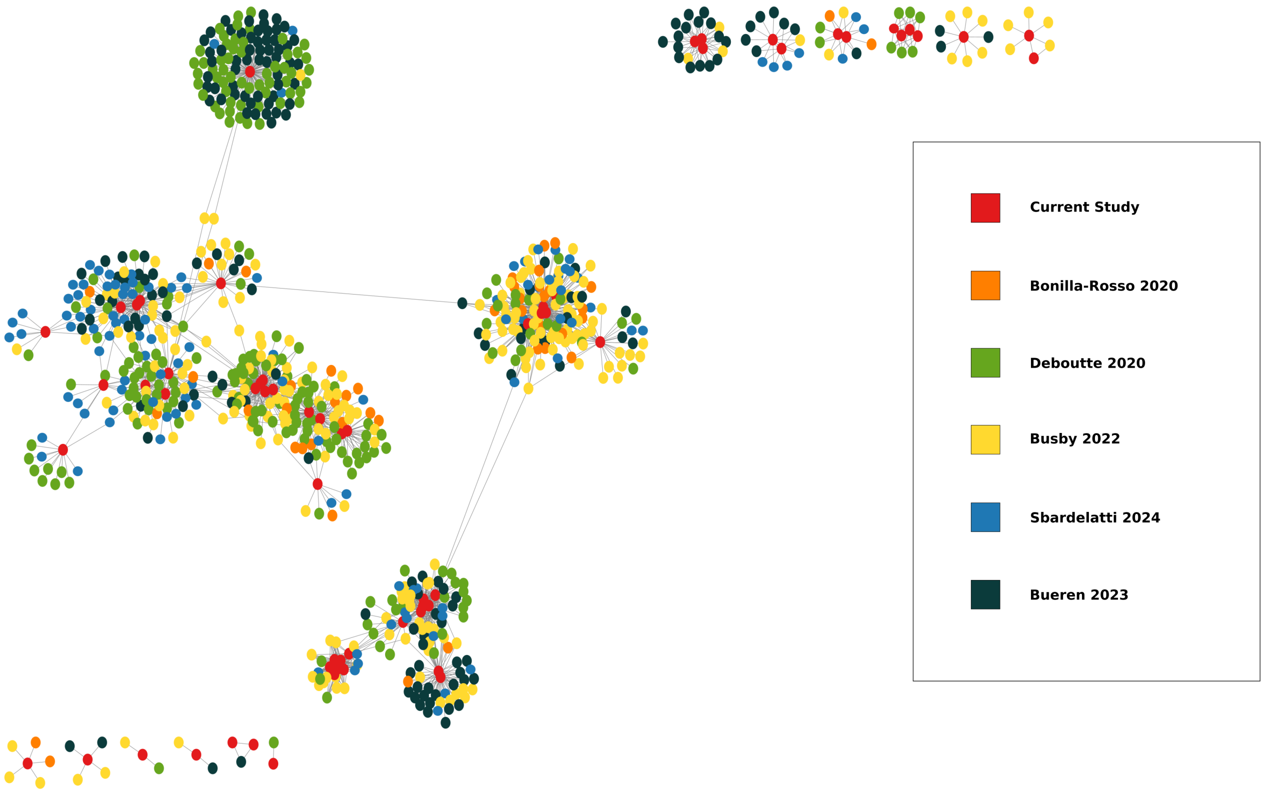


Supplemental Figure 1: vConTACT3 network generated from honey bee phage genomes recovered from this study (red). The remaining nodes are colored by study that they were recovered from.

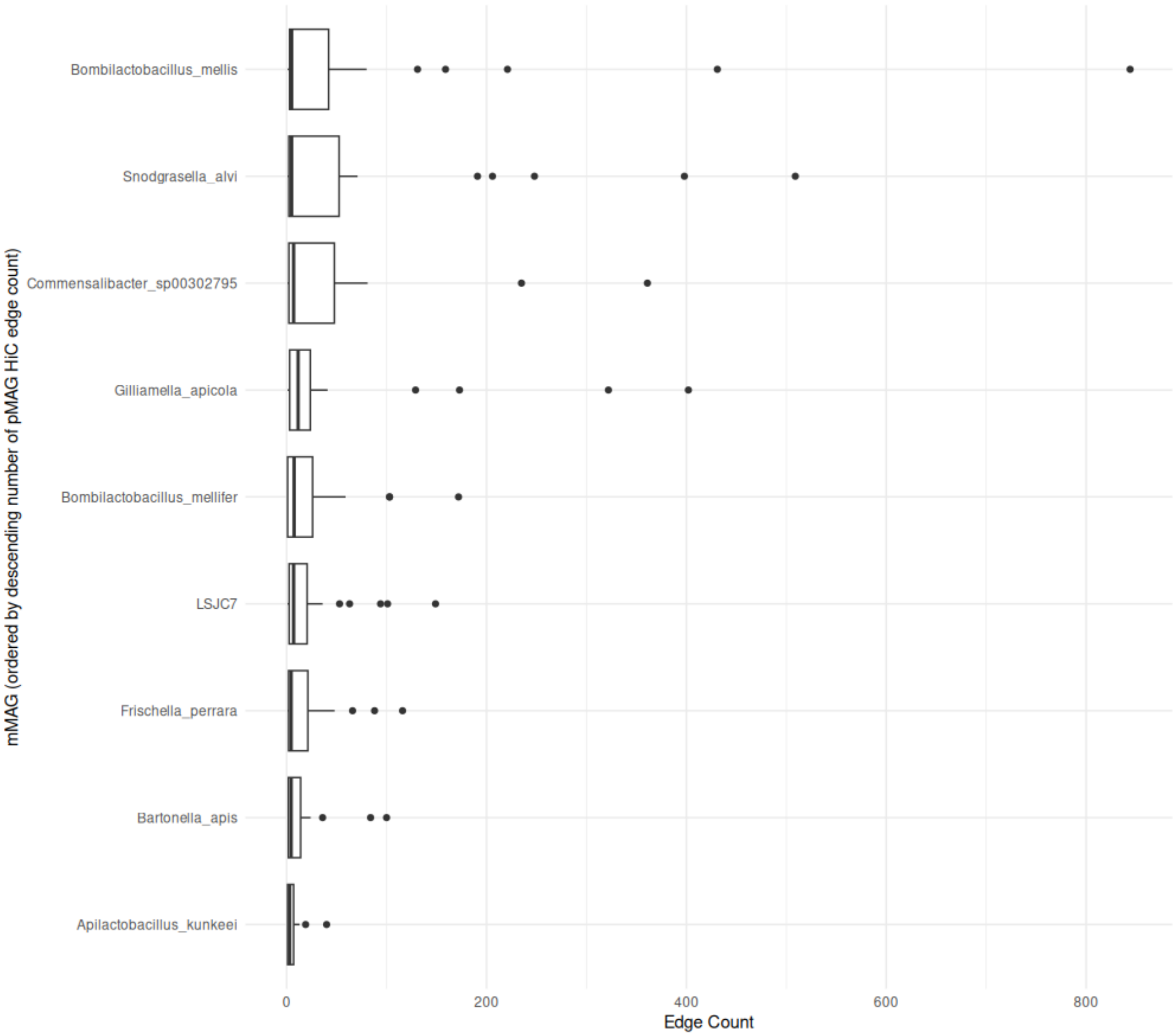


Supplemental Figure 2: HiC edge count distribution for all mMAG x pMAG interactions. Edge count are summed across all metagenomes.

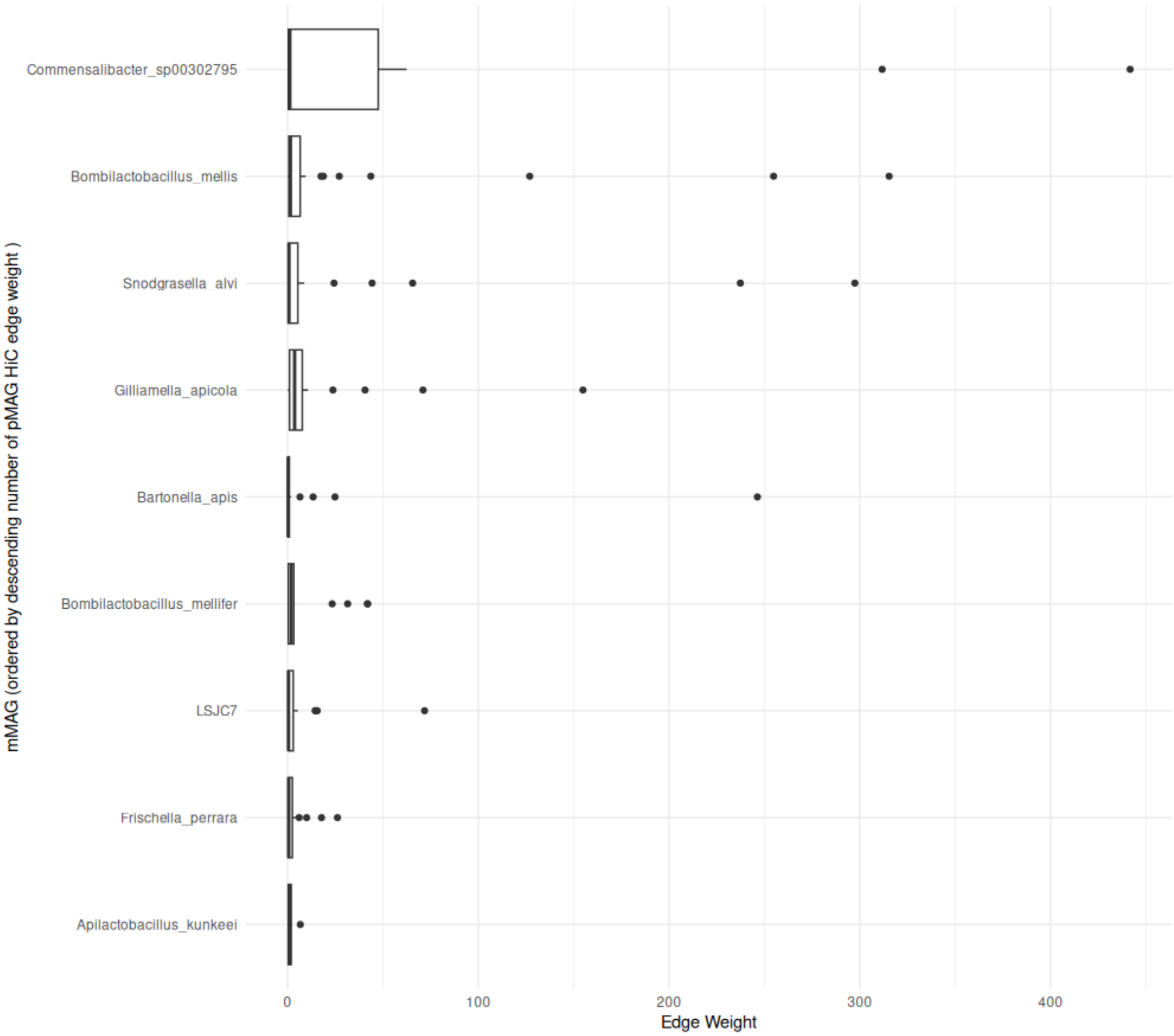


Supplemental Figure 3: HiC edge weight distribution for all mMAG x pMAG interactions. Edge weights are summed across all metagenomes.

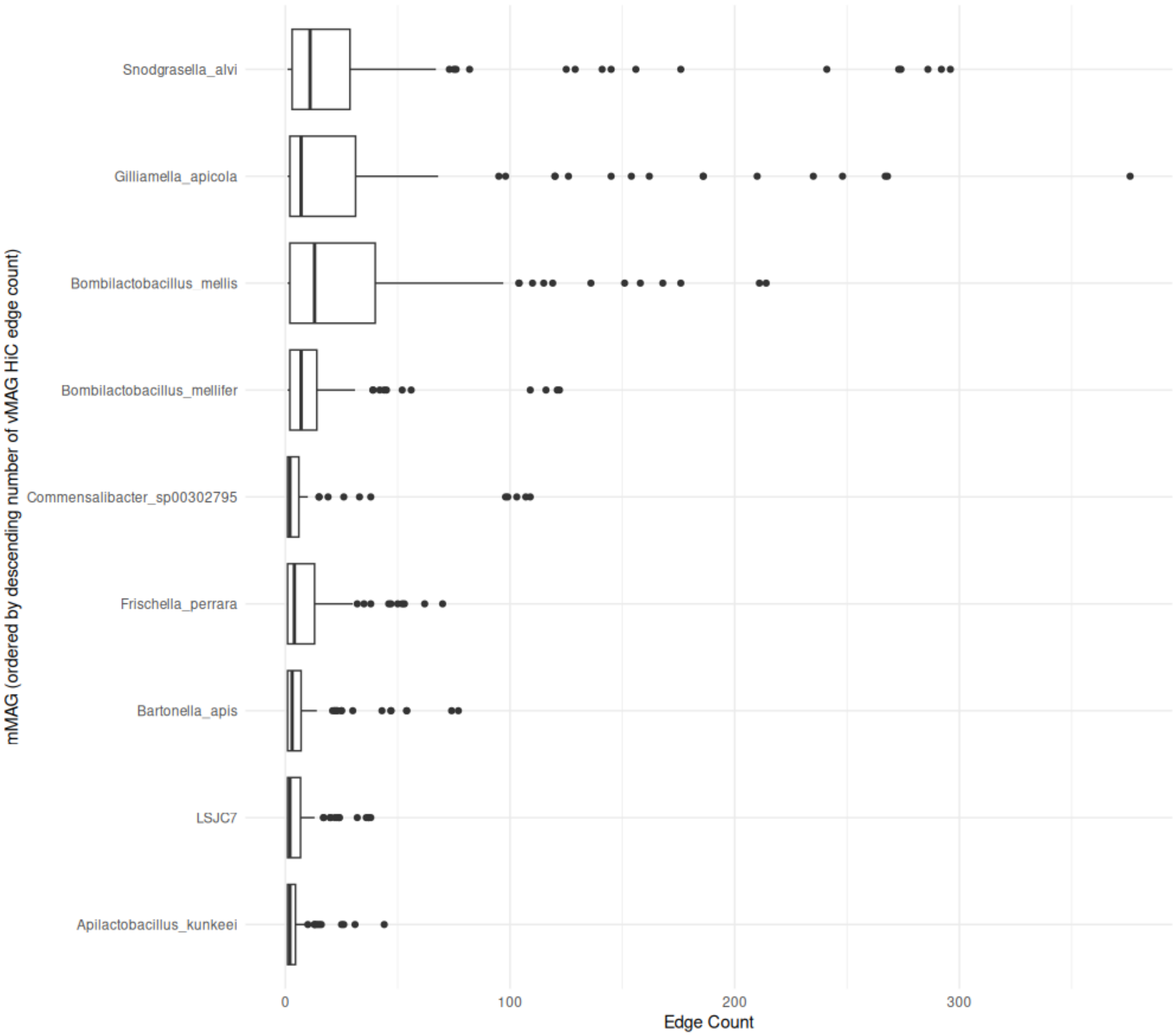


Supplemental Figure 4: HiC count distribution for all mMAG x vMAG interactions. Edge counts are summed across all metagenomes.

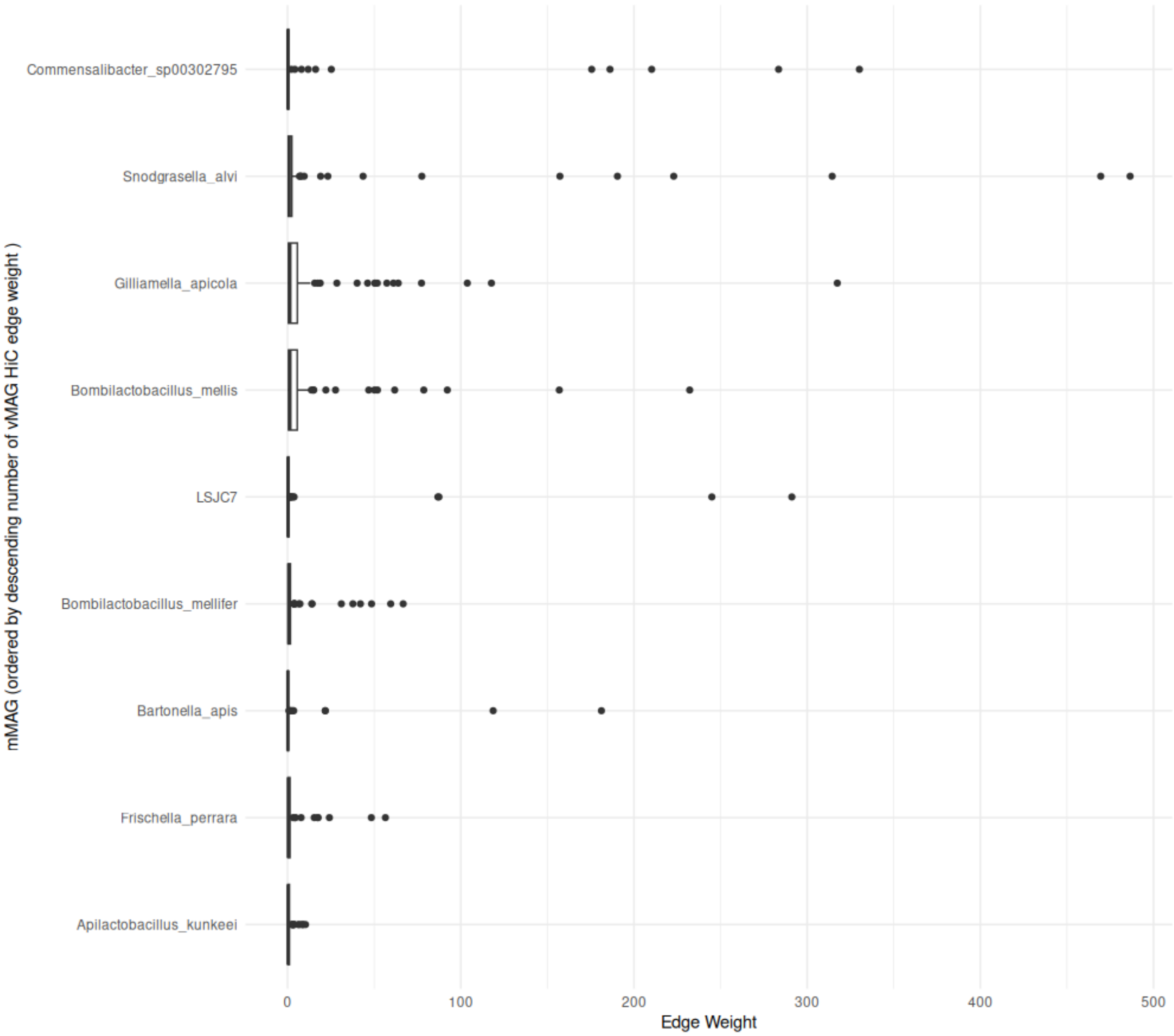


Supplemental Figure 5: HiC edge weight distribution for all mMAG x vMAG interactions. Edge counts are summed across all metagenomes.

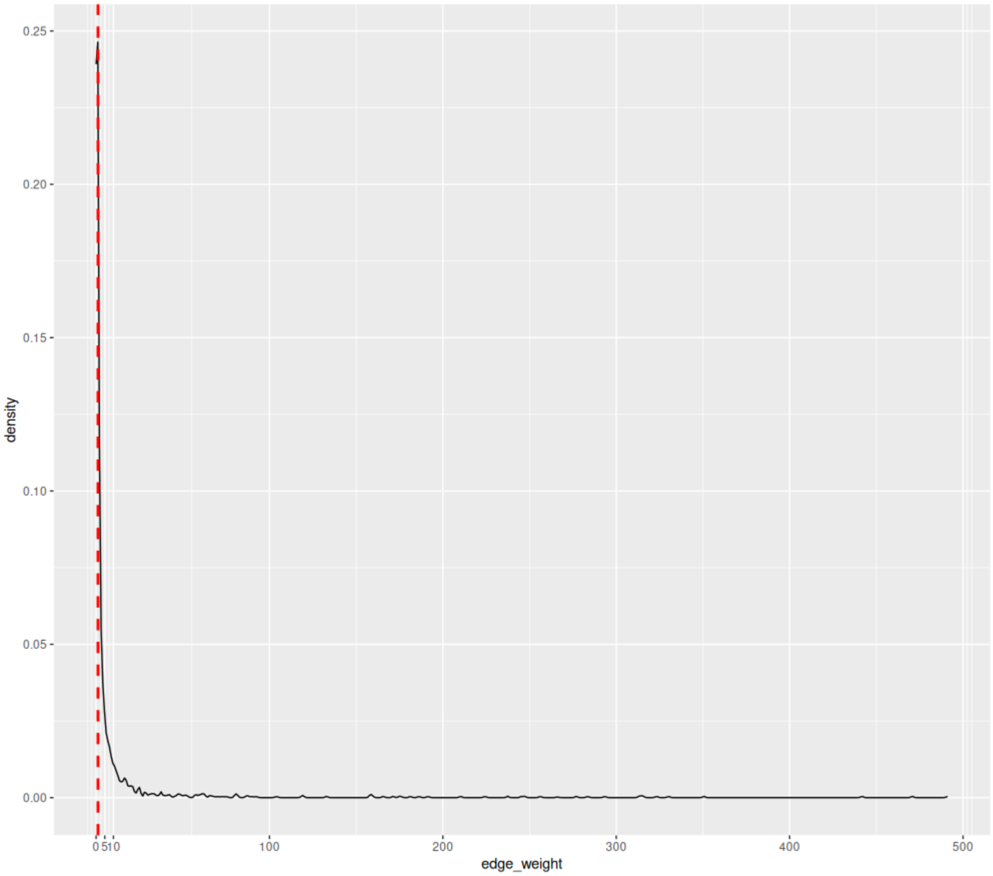


Supplemental Figure 6: Edge weight distribution. Dotted, red vertical line shows average edge weight for *A. kunkeei*.

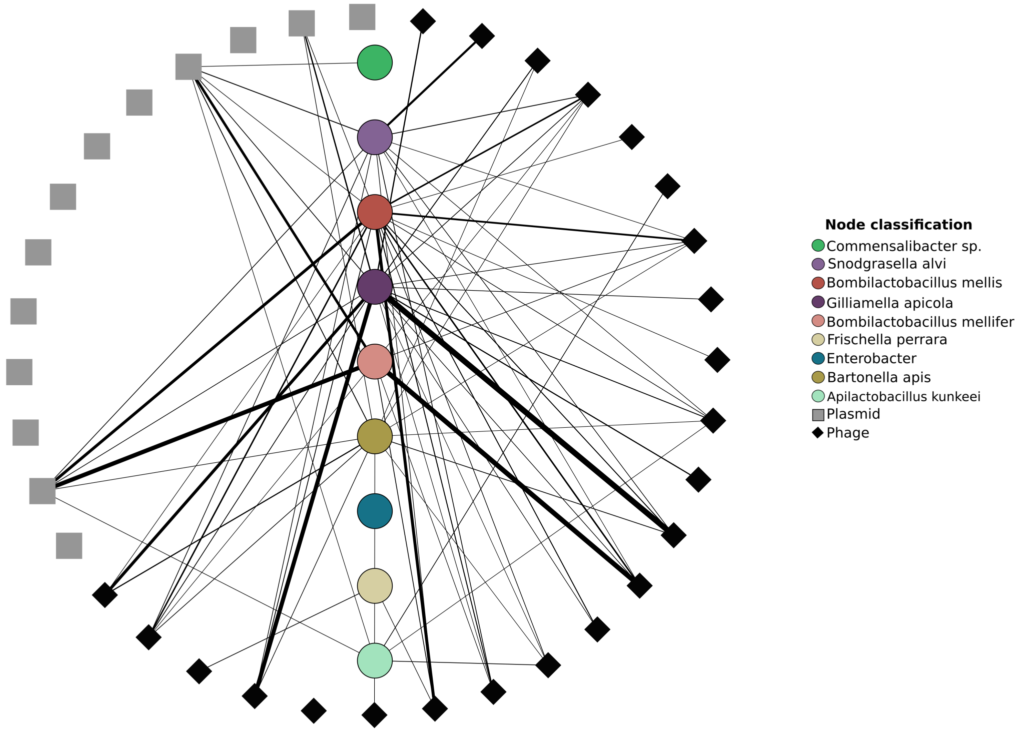


Supplemental Figure 7. Individual HiC network for Metagenome A.

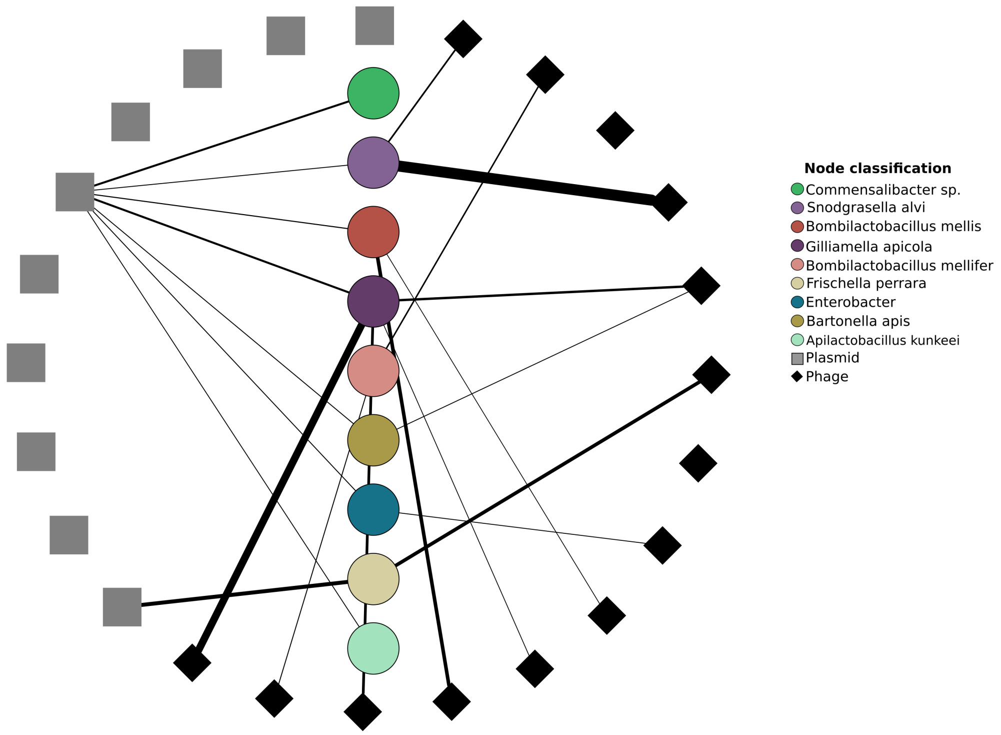


Supplemental Figure 8. Individual HiC network for Metagenome B.

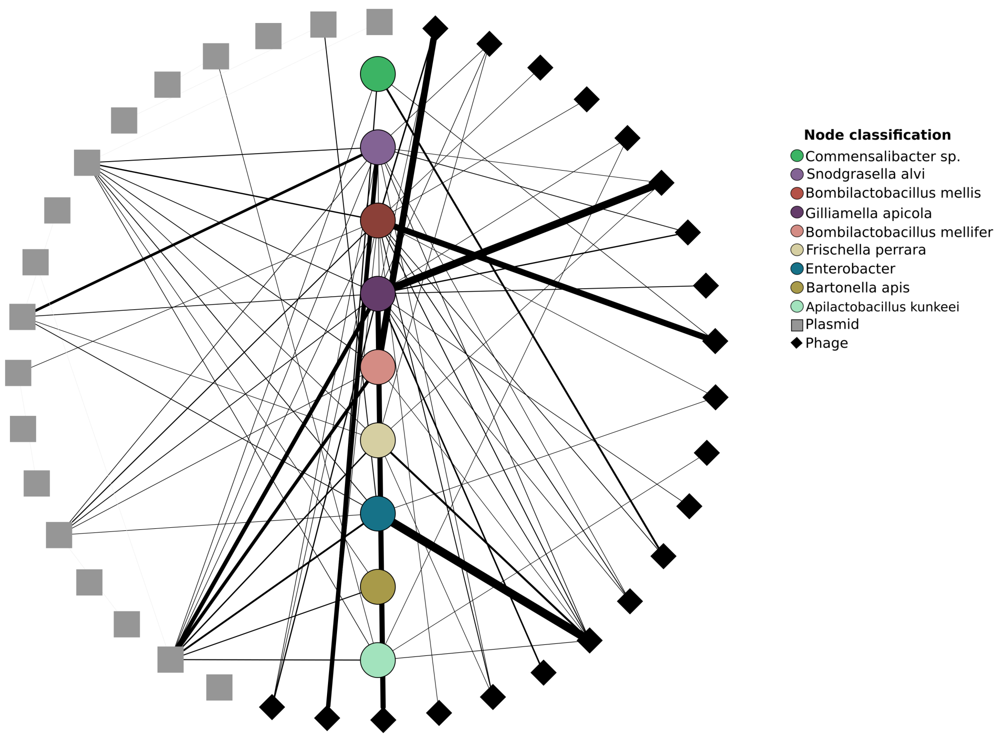


Supplemental Figure 9. Individual HiC network for Metagenome C.

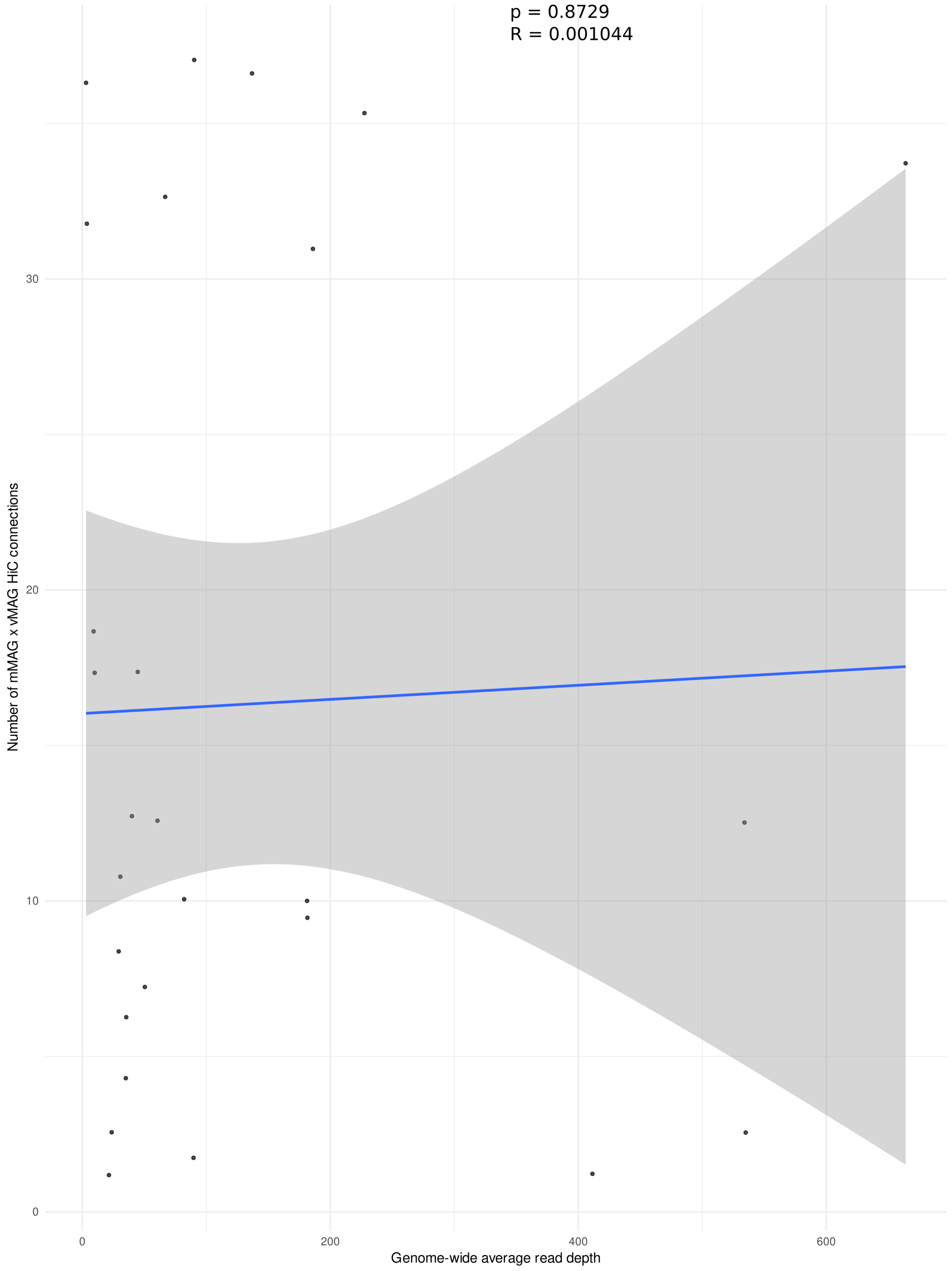


Supplemental Figure 10. X-axis shows the genome-wide average read depth for mMAGs while the Y-axis shows the number of mMAG x pMAG HiC connections. Blue line is generated from "lm" from geom_smooth. Shaded area represents 95% CI.

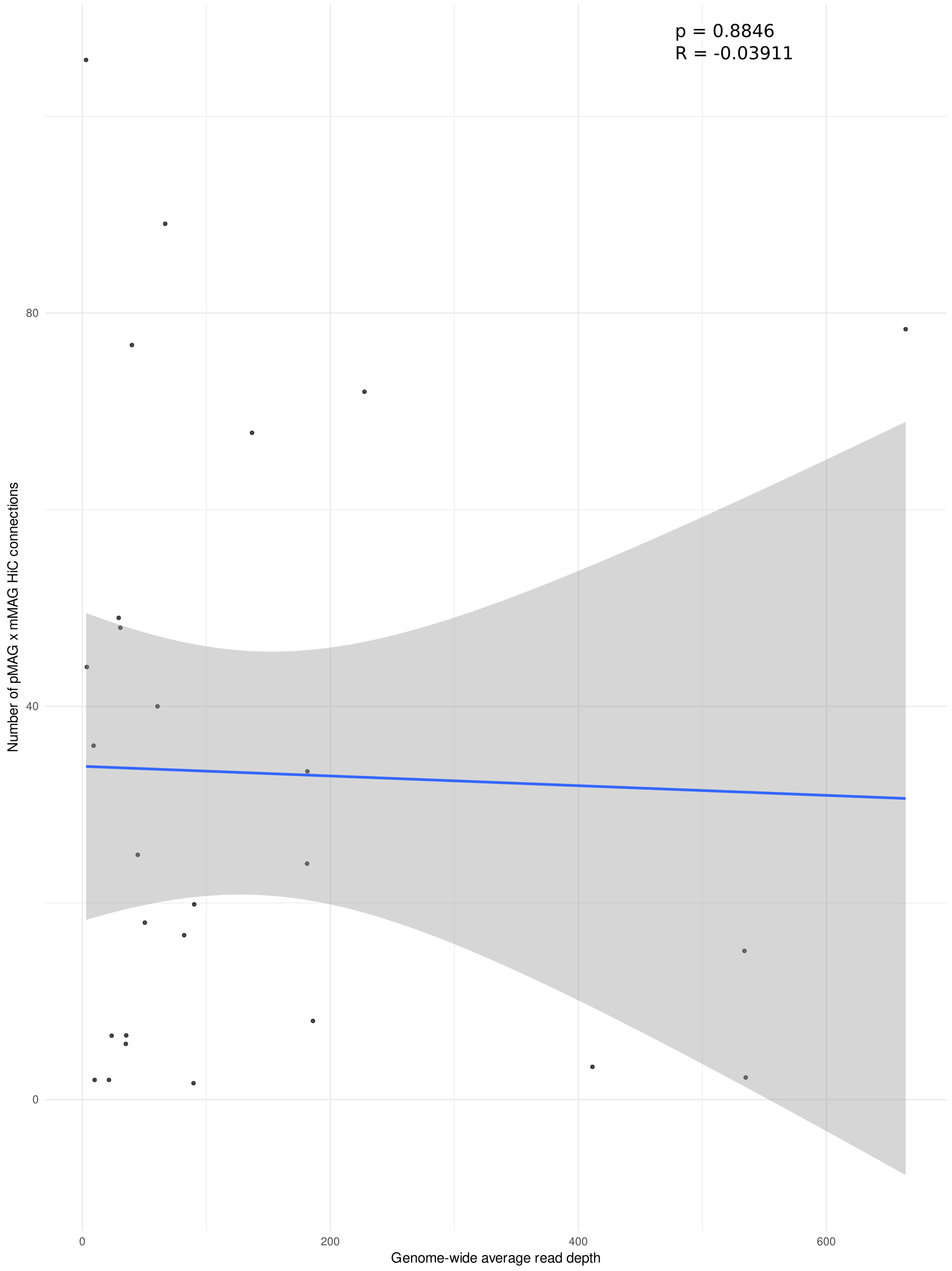


Supplemental Figure 11. X-axis shows the genome-wide average read depth for mMAGs while the Y-axis shows the number of mMAG x vMAG HiC connections. Blue line is generated from "lm" from geom_smooth. Shaded area represents 95% CI.

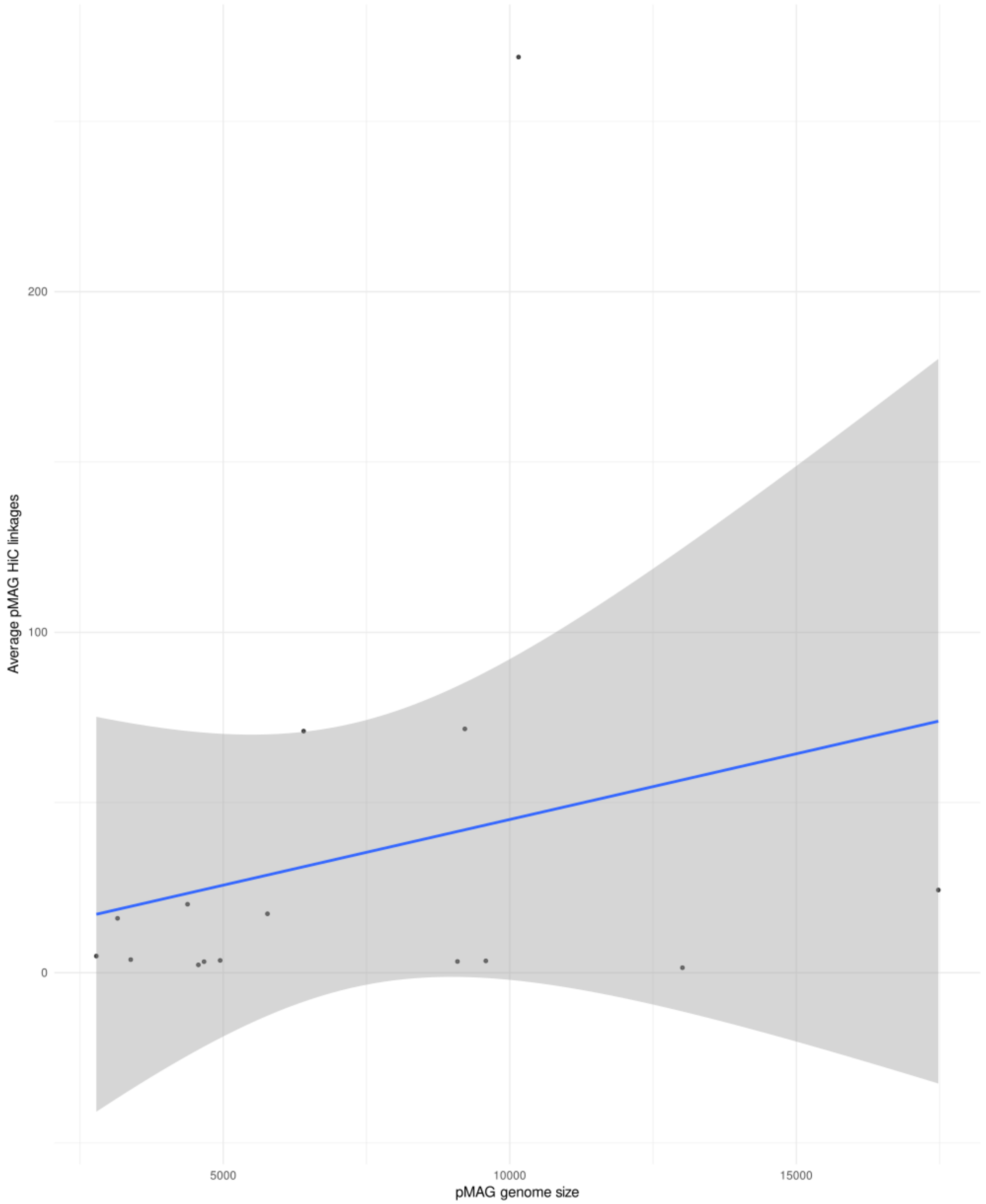


Supplemental Figure 12. X-axis shows the genome size for pMAGs while the Y-axis shows the average number of mMAG x pMAG HiC connections. Blue line is generated from "lm" from geom_smooth. Shaded area represents 95% CI.

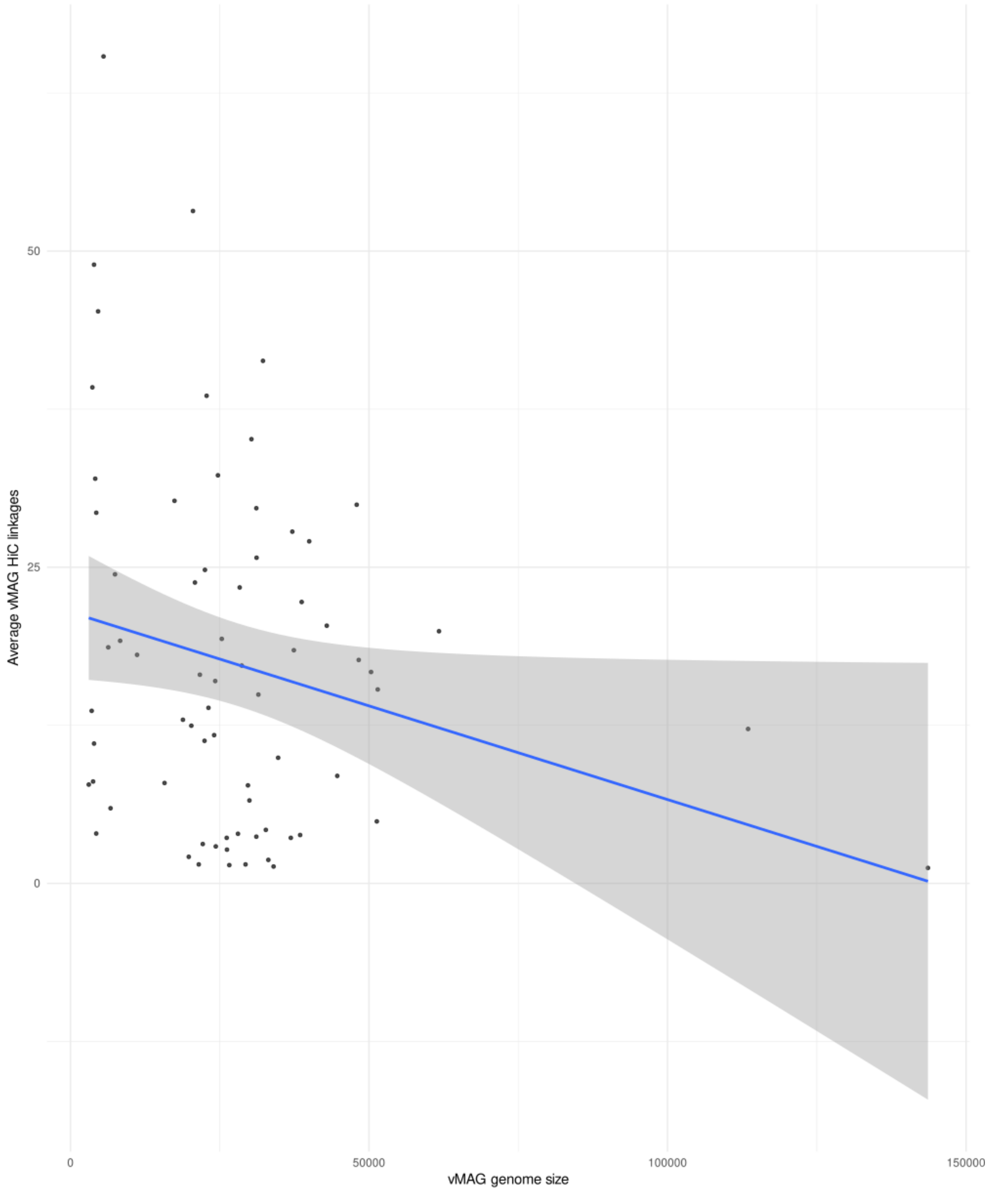


Supplemental Figure 13. X-axis shows the genome size for vMAGs while the Y-axis shows the average number of mMAG x vMAG HiC connections. Blue line is generated from ”lm” from geom_smooth. Shaded area represents 95% CI.

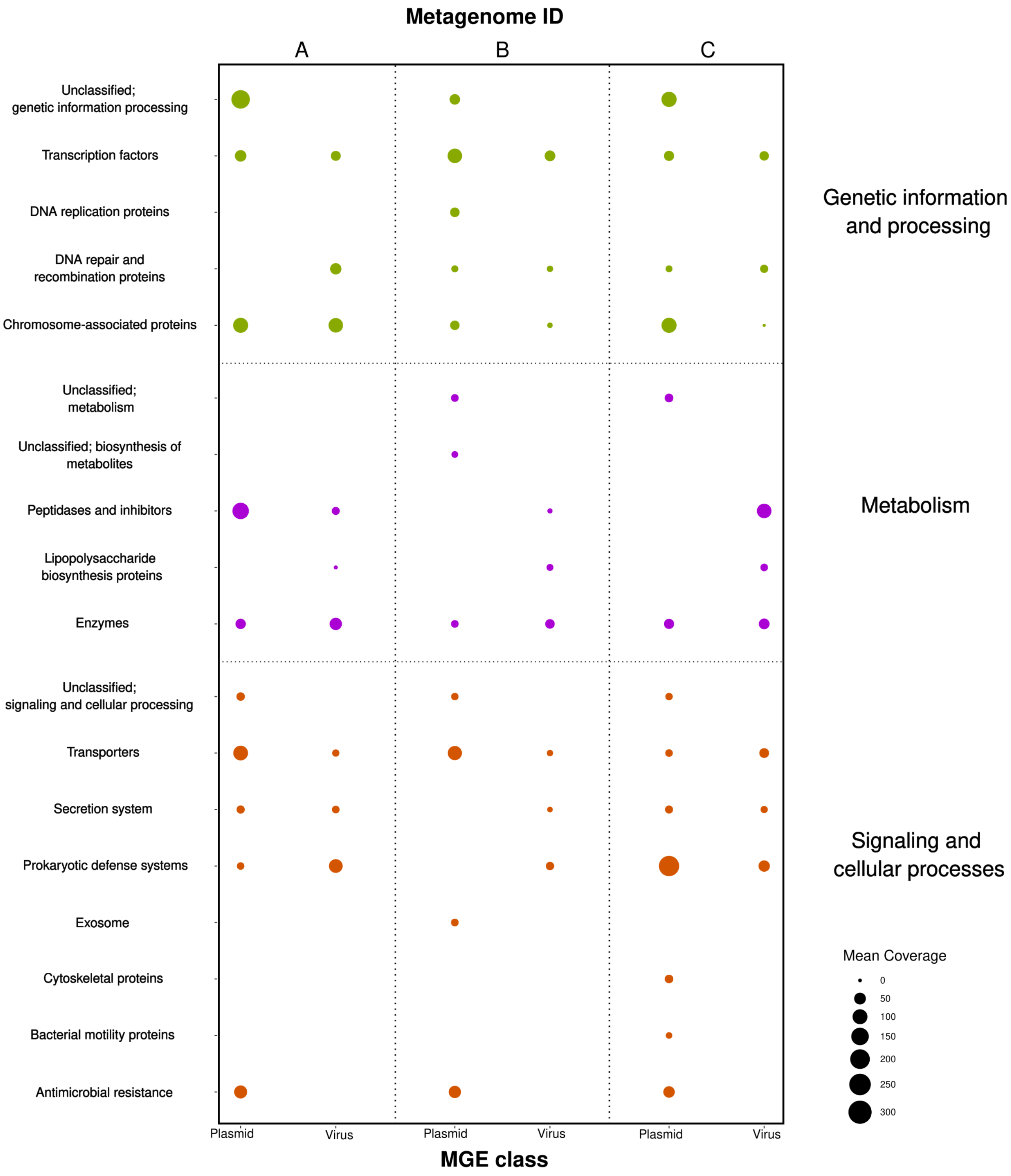


Supplemental Figure 14. Mean coverage for genes associated with either vMAGs or pMAGs (bottom x-axis) within each metagenome (top x-axis, separated by dotted line). Genes were grouped by association with the KEGG BRITE database (y-axis). Broad level groupings are demarcated by color and increased dot size corresponds to higher overall mean coverage for the corresponding MGE-KEGG BRITE grouping.

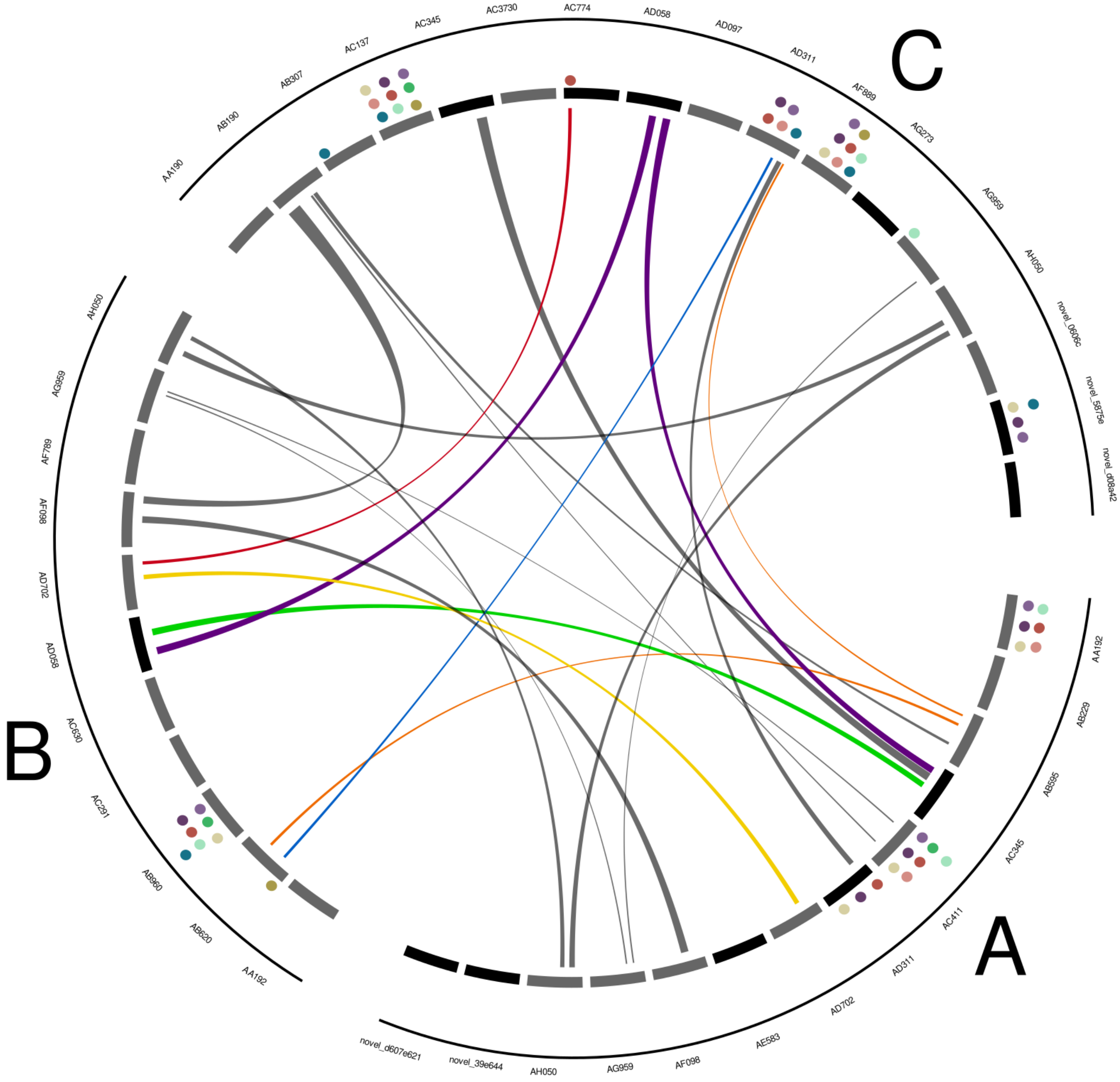


Supplemental Figure 15. Circos plot from Figure 3B. Individual plasmid IDs are given above each plasmid.

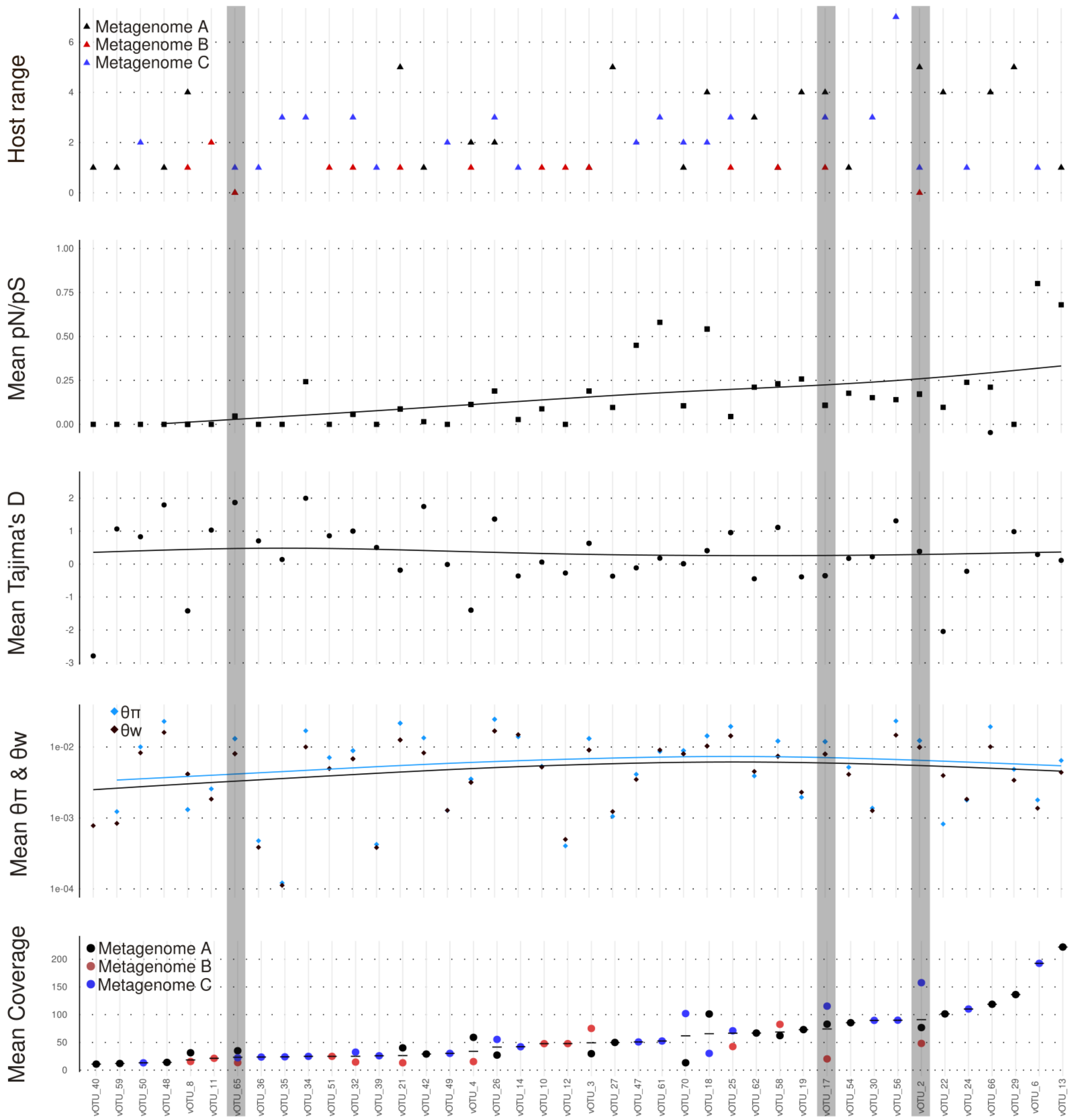


Supplemental Figure 16. Genomic variation statistics and host range calculated for vMAGs with at least 1 post-filtered HiC connection with an mMAG host (x-axis). vMAGs that appear in all 3 metagenomes are highlighted in gray. Mean metagenomic coverage for each vMAG is shown at the bottom plot (colored dots); if a vMAG was associated with ≥2 metagenomes, the average coverage is depicted as a solid black line. Genome-wide averages for nucleotide diversity (blue diamond) and $\theta$w(black diamond), Tajima's D (black dots, third plot), and pN/pS (black squares, fourth plot) are largely stable with respect to coverage. Individual host ranges associated vMAGs are shown at top of plot (colored triangles).

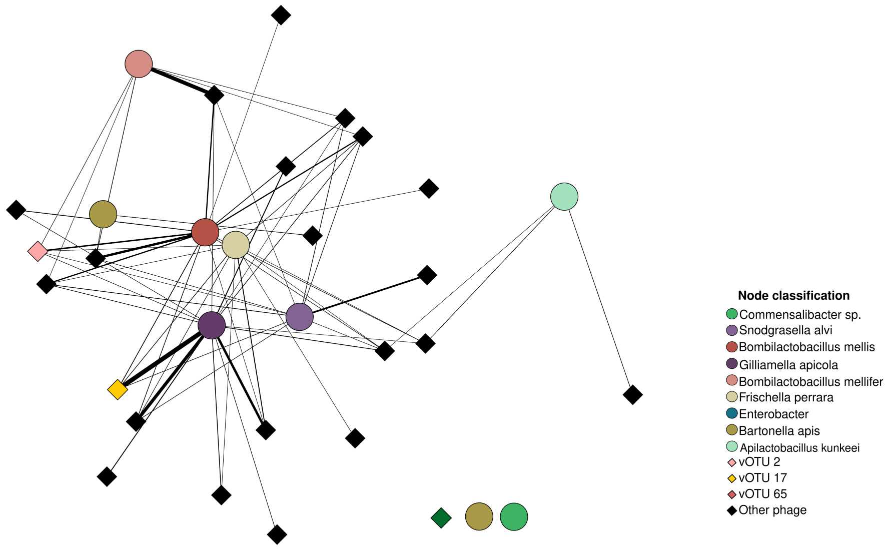


Supplemental Figure 17. Viral network for metagenome A. Highlighted diamonds (vOTU 2, vOTU 17, and vOTU 65) represent phages that are shared between all 3 metagenomes.

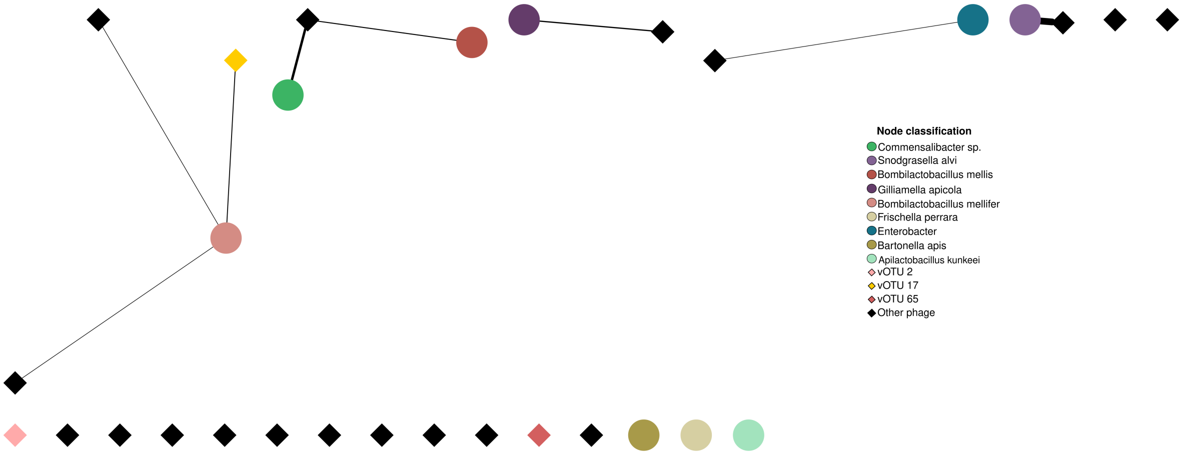


Supplemental Figure 18. Viral network for metagenome B. Highlighted diamonds (vOTU 2, vOTU 17, and vOTU 65) represent phages that are shared between all 3 metagenomes.

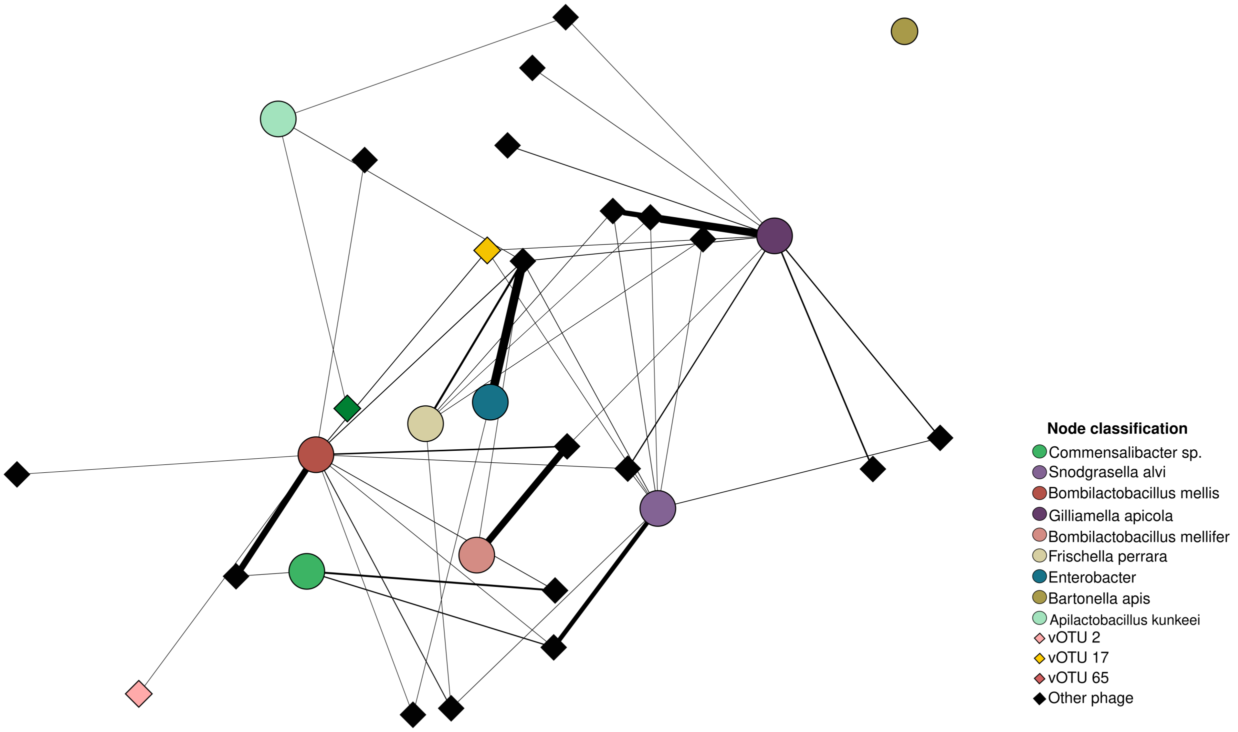


Supplemental Figure 19. Viral network for metagenome C. Highlighted diamonds (vOTU 2, vOTU 17, and vOTU 65) represent phages that are shared between all 3 metagenomes.

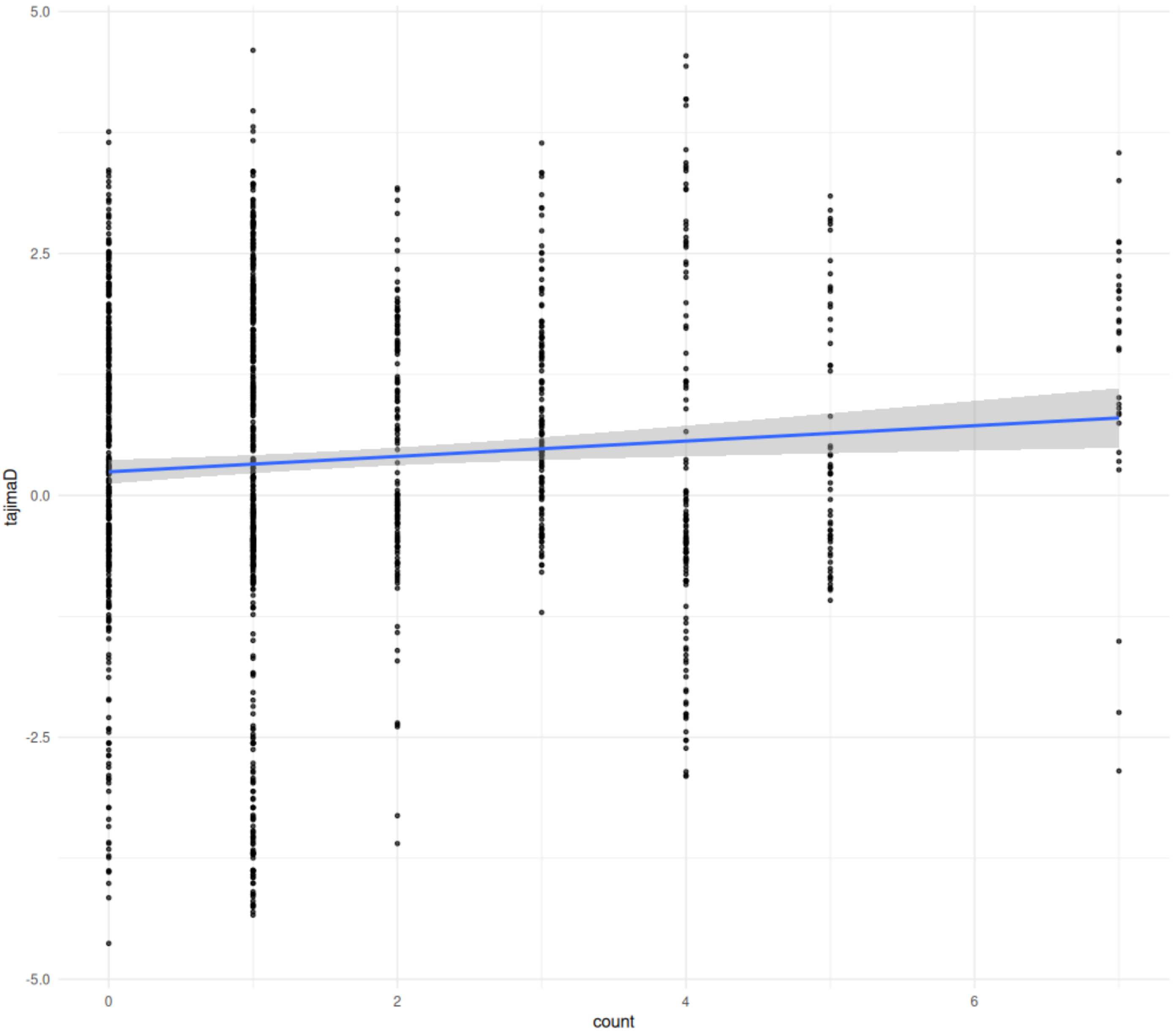


Figure 1: Supplemental Figure 20. Correlations between the number of intrametagenomic mMAG x vMAG filtered HiC contacts (x-axis) and genic measures of *Tajima's D*. Regression line generated from geom_smooth (method=lm) and shaded area represents 95% CI.
